# UniCure: A Foundation Model for Predicting Personalized Cancer Therapy Response

**DOI:** 10.1101/2025.06.14.658531

**Authors:** Zexi Chen, Saisai Tian, Jiazheng Pei, Ruichu Gu, Yongge Li, Shizhi Ding, Yaqian Xu, Xinlong Zheng, Miaoyu Liu, XinXing Du, Junchao Zhu, Jiawei Zou, Jing Xu, Wenli Jiang, Chen Ye, Baijun Dong, Qi Zhang, Shengxiang Ren, Shu Wang, Han Wen, Weidong Zhang, Luonan Chen

## Abstract

Predicting drug efficacy across diverse patient contexts remains a major challenge in oncology, as models trained on cancer cell lines often fail to capture patient-specific biology. Emerging biological foundation models and patient-derived technologies offer a promising solution. Here, we present UniCure, the first pre-trained foundation model integrating both omics and chemical foundation models (UCE and Uni-mol) to predict transcriptomic responses to drugs across diverse cellular and tissue contexts, enabling personalized cancer therapy and drug prioritization at the individual level. Rather than encoder/decoder used in traditional models, UniCure utilizes parameter-efficient fine-tuning (PEFT) techniques for optimizing the training process, a novel FlexPert module for modeling flexible drug-cell interactions, and a Maximum Mean Discrepancy (MMD) loss for learning unpaired data. Trained on over 1.8 million perturbation RNA-seq profiles over 22,000 compounds, 166 cell types, and 24 tissues, UniCure achieves high accuracy in predicting both dose-dependent responses and drug combination effects, demonstrating strong generalization across bulk and single-cell transcriptomic data. In particular, fine-tuning on our patient-derived tumor-like clusters and real-world data of 800 profiles enables UniCure to generate individualized therapeutic predictions on patient’s tissue samples. In addition, UniCure enables patient stratification based on the predicted drug responses, providing a new way for subtyping patients. UniCure’s drug prioritization was validated across over 1000 patients from pan-cancer cohorts and supported by experiments of candidate therapeutics. By enabling the potential to screen millions of compounds per patient at scale, UniCure represents a biologically grounded tool that could advance personalized precision oncology and accelerate drug discovery.

## Main

Accurately predicting the complex biological effects of drugs across cell types and individuals could markedly improve clinical translation success and facilitate personalized therapies, particularly for patients with refractory advanced cancers.

Leveraging high-throughput sequencing technologies, such as next-generation sequencing and single-cell RNA sequencing, we can now decipher the intricate molecular mechanisms underlying drug responses with unprecedented resolution, particularly at the transcriptomic level. Vast amounts of drug-related transcriptomic data, generated through these advancements and publicly available in databases like Connectivity Map (CMap)^1^, Integrated Network-based Cell-Signature (LINCS)^2^, sci-Plex^3^, and Tahoe-100M^4^, provide invaluable resources for drug discovery and understanding the complexities of drug action. Despite their utility, cell line models demonstrate limited capacity to recapitulate clinical tumor biology, constrained by insufficient modeling of tumor microenvironment complexity, intratumoral heterogeneity, and patient-specific pharmacodynamic determinants^5–7^. Therefore, the application of encoder-decoder models like DLEPS^8^, DeepCE^9^, ChemCPA^10^, TransiGen^11^, and PRnet^12^ to personalized medicine is limited by their reliance on public cell line data and their inability to model complex patient-specific drug responses.

To bridge this gap and harness the full potential of these rich datasets, the rise of foundation models in biology offers a promising avenue. These models, such as Geneformer^13^, scGPT^14^, scFoundation^15^, and GeneCompass^16^, are pre-trained on massive datasets of diverse biological contexts, endowing them with a broad understanding of cellular function and enabling more robust and generalizable representations of cell states. Notably, Universal Cell Embedding (UCE) integrates protein structural data via ESM2 embeddings, offering a framework with enhanced potential for drug response modeling by capturing multimodal cellular states^17^. To efficiently adapt models for specific tasks, Parameter-Efficient Fine-tuning (PEFT) techniques such as Low-Rank Adaptation (LoRA)^18^, Prompt Tuning^19^, and Prefix Tuning^20^ modify only a minimal set of parameters, preserving the pre-trained knowledge while avoiding catastrophic forgetting. By leveraging these powerful (or fine-tuned) foundation models, it becomes feasible to extend the applicability of cell line-derived data to individual patient scenarios. Concurrently, recognizing the limitations of traditional cell lines, several technologies have emerged that better preserve the unique characteristics of individual patient tumors and enable more accurate predictions of drug efficacy, including DNA sequencing for targeted therapy, patient-derived tumor xenografts (PDXs)^21^, patient-derived organoids (PDOs)^22^, and patient-derived tumor-like cell clusters (PTCs)^23, 24^. Integrating the generalizable insights obtained from foundation models with the patient-specific context captured by these technologies holds the key to unlocking truly personalized drug response predictions.

Here, we introduce UniCure, the first pre-trained multi-modal foundation model that integrates both omics and chemical foundation models to predict transcriptomic responses to drugs across diverse cellular and tissue contexts, enabling personalized cancer therapy and drug prioritization at the individual level. Rather than encoder/decoder used in traditional models, UniCure leverages pre-trained foundation models, namely Uni-mol^25^ for drug representation and UCE for cell representation, with Low-Rank Adaptation (LoRA)^18^ fine-tuning, significantly enhancing the model’s generalization capacity and efficiency. Notably, it features a unique FlexPert module that can process flexible drug inputs through a sliding window module and effectively model complex drug-cell interactions through a cross-attention mechanism and a biologically-inspired staged-training strategy. Additionally, UniCure addresses the challenge of unpaired pre- and post-perturbation data using Maximum Mean Discrepancy (MMD)^26^, which enables the model to learn the distributional shift induced by drug treatment without requiring paired samples. Furthermore, UniCure inherits vast data information of 209 million molecular structures and 36 million cell data points, and is further trained on massive data including over 1.8 million cell line and single-cell perturbation data points across over 22,000 compounds, 166 cell types, and 24 tissues. Crucially, it is then fine-tuned on our approximately 800 patient-derived drug perturbation datasets, encompassing both PTCs (TNBC, LUAD, and BLCA) and real-world data (Hematologic Malignancies, Breast Cancers, and Ovarian Cancers), to capture patient-specific responses and tumor microenvironment complexities.

Through extensive evaluations detailed in the subsequent sections, we demonstrate that UniCure achieves high fidelity in predicting transcriptomic perturbations of drugs across diverse public datasets, significantly outperforming existing approaches. Unlike previous models focusing solely on transcriptomic drug perturbation prediction, UniCure introduces the following several advancements that expand its capabilities.

1. First, leveraging LoRA fine-tuning, UniCure unifies representations for cell lines derived from different platforms, effectively addressing batch effects.
2. Second, for the first time, UniCure integrated cell representation foundation model UCE and chemical foundation model Uni-Mol, demonstrated pronounced capability and generalizability in drug efficacy and combination predictions.
3. Third, UniCure achieves robust, individualized predictions through efficient fine-tuning on our 800 profiles comprised of patient-derived tumor-like clusters (PTCs) and real-world data, demonstrating high correlation even with this limited dataset.

Applied to over 1000 patients from public clinical cohorts, UniCure enables novel patient stratification based on the predicted therapy response profiles, consistent with prior findings^27^. Moreover, these personalized rankings of administered therapies, validated from patients of clinical cohorts, not only correlate significantly with overall survival but also prioritize known targeted therapies within their indicated cancer types, further supporting the framework’s translational potential. Building on these capabilities, UniCure further identified promising natural products against triple-negative breast cancer, lung adenocarcinoma, and bladder cancer, with *in vitro* assays confirming their potent, concentration-dependent inhibition of cancer cell viability and colony formation. Collectively, these results illuminate UniCure as a versatile foundation model for drug discovery and personalized oncology, bridging foundational research and clinical application.

## Result

### Overview of UniCure model

We introduce UniCure, the first pre-trained foundation model integrating both omics and chemical foundation models (UCE and Uni-mol) to predict transcriptomic drug responses across diverse cell types and tissues, enabling personalized cancer therapy and drug prioritization at the individual level. It aims to bridge the gap between traditional drug discovery and personalized medicine by accurately modeling compound-cell interactions and enabling patient-specific drug response prediction.

UniCure addresses the problem of learning a function, 𝑓(⋅), parameterized by model weights 𝜃, that maps a cell’s initial transcriptomic state, 𝑋 ∈ ℝ^𝑛^ (where 𝑛 represents the number of genes), and a set of perturbations, 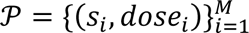 (where 𝑠_𝑖_ represents 𝑖 drug’s SMILES strings and *dose* represents the dose of the drug), to the perturbed transcriptomic state 𝑋′_𝑖_ ≈ 𝑓(𝑋_𝑖_, 𝒫_𝑖_; 𝜃).

To achieve this, UniCure leverages pre-trained foundation models for molecular and cellular representations (Fig. 1a). It utilizes Uni-mol to encode drug molecules into embedding vectors and UCE with a LoRA module to represent cell states (Methods). Specifically, a novel FlexPert module, leveraging sliding window-enhanced cross-attention mechanisms, effectively captures the complex interactions between flexible drug inputs and diverse cellular contexts (Methods and Supplementary Fig. 1a). Furthermore, a distribution-based loss function, Maximum Mean Discrepancy (MMD)^28^, addresses challenges arising from unpaired pre- and post-perturbation data (Methods). In contrast to Gaussian negative log-likelihood losses and optimal transport-based methods^12, 29^, MMD enables the model to learn complex relationships without requiring paired data in every instance and, critically, exhibits improved generalization to unseen drugs.

**Fig. 1:**
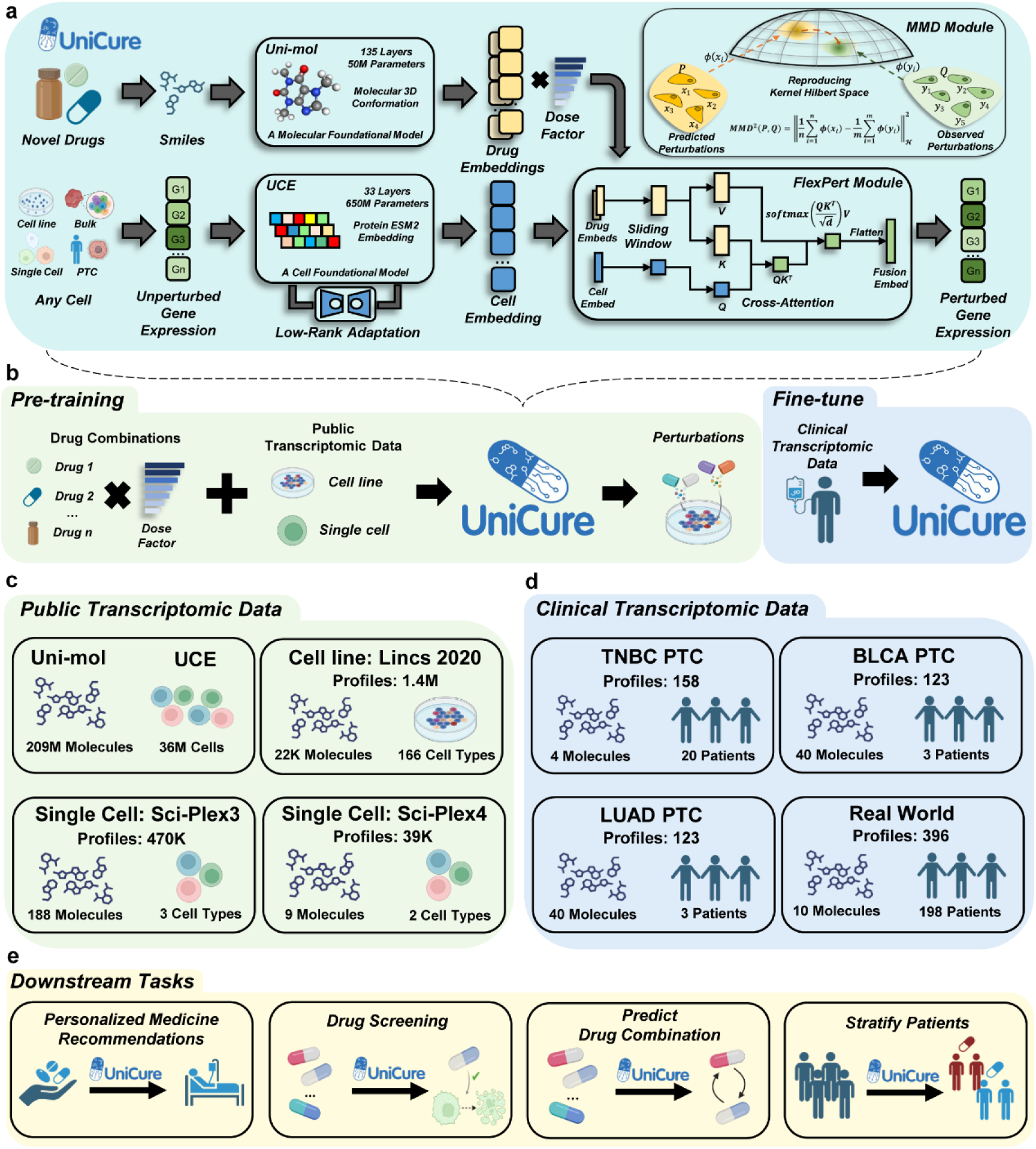
Overview of the UniCure model. **a**, UniCure is the first foundation model integrating both omics and chemical foundation models (UCE and Uni-mol) to predict drug-induced transcriptomic perturbations across diverse human cellular and tissue systems, built on four core modules: foundation models with LoRA, FlexPert module, MMD-based alignment, and staged training. **b**, Training strategy of UniCure involves pre-training on multi-drug perturbations across cell lines and single-cell data, considering drug dose effects, followed by fine-tuning with clinical PTC samples for personalized predictions. **c**, UniCure is built on extensive public data, leveraging Uni-mol (209M molecular conformations) and UCE (36M cells), and is further trained on perturbation profiles from cell lines (LINCS 1000, 1.4M profiles) and single-cell datasets (Sci-Plex3, 470K profiles; Sci-Plex4, 39K profiles). **d**, UniCure is fine-tuned and validated using clinical data, including TNBC PTC (79 profiles, 20 samples, 4 drugs), BLCA PTC (123 profiles, 3 samples, 40 drugs), LUAD PTC (123 profiles, 3 samples, 40 drugs), and 396 real-world patient-derived perturbation profiles covering hematologic malignancies, breast cancers, and ovarian cancers. **e,** UniCure enables diverse downstream tasks, including clinical personalized medicine recommendations, drug screening and optimization, combination therapy prediction, drug response prediction, and patient stratification.

UniCure’s architecture and training strategy confer several key advantages. The use of pre-trained models, coupled with LoRA fine-tuning, mitigates catastrophic forgetting during adaptation, preserving pre-trained biological knowledge while enabling effective transfer learning across diverse cell types and compound classes^18^. UniCure is trained using a staged strategy to enhance computational efficiency and functional modularity (Methods and Supplementary Fig. 1b). The first stage, focused on stabilizing cell environment, utilizes the vast amount of information contained within the UCE with LoRA, query projection layer, and UniCure decoder to build a reliable baseline to test the impact of various perturbations. The second stage leverages a key-value projection layer and FlexPert decoder in order to accurately predict post-perturbed states. By leveraging an MMD loss and transmission cost constraint, the model is able to produce accurate and reliable predictions regarding the state of cellular function (Methods).

Enabled by its modular architecture, UniCure was pre-trained on a wide variety of drug combinations, dosages, cell lines, and single-cell data, and further refined with real-world and PTCs data (Fig. 1b). As a foundation model framework integrating Uni-mol (representing 209M molecular conformations) and UCE (encompassing 36M cells) (Fig. 1c), UniCure was further trained on a vast collection of perturbation data, encompassing cell line datasets (LINCS 2020) and large-scale single-cell datasets (Sci-Plex3, and Sci-Plex4), totaling over 1.8 million perturbation data points across over 22,000 compounds, 166 cell types, and 24 tissues. Critically, UniCure was then further fine-tuned on our approximately 800 patient-derived drug perturbation samples, including PTCs and real-world data (Fig. 1d), to capture patient-specific responses and the complexities of the tumor microenvironment. This allows UniCure to be rapidly adapted to individual patients, thereby enabling accurate prediction of drug responses at the individual patient’s transcriptome level.

This framework enables a diverse range of downstream tasks (Fig. 1e), including drug screening, personalized medicine recommendations, combination therapy prediction, and response-based patient stratification. The subsequent sections will detail the empirical evaluation of UniCure’s performance in these areas.

### UniCure Accurately Predicts Transcriptomic Perturbations by Learning Robust Cellular Representations

UniCure was trained on diverse large-scale perturbation datasets, during which it exhibited a stable decline in both training and validation loss, alongside consistent improvements in key evaluation metrics—demonstrating the robustness of the learning process (Fig. 2a and Extended Data Fig. 1a-c). We then evaluated the model’s predictive accuracy on held-out subsets from each dataset. On LINCS2020, UniCure achieved strong agreement between predicted and observed landmark gene expression changes, with Pearson and Spearman correlation coefficients both exceeding 0.9 and an R² above 0.8. Similarly high performance was observed on single-cell datasets: for SciPlex3, Pearson correlation surpassed 0.9 and Spearman exceeded 0.8; for SciPlex4, which includes dual-compound perturbations, Pearson correlation remained above 0.9, and Spearman correlation exceeded 0.8 (Fig. 2b).

**Fig. 2:**
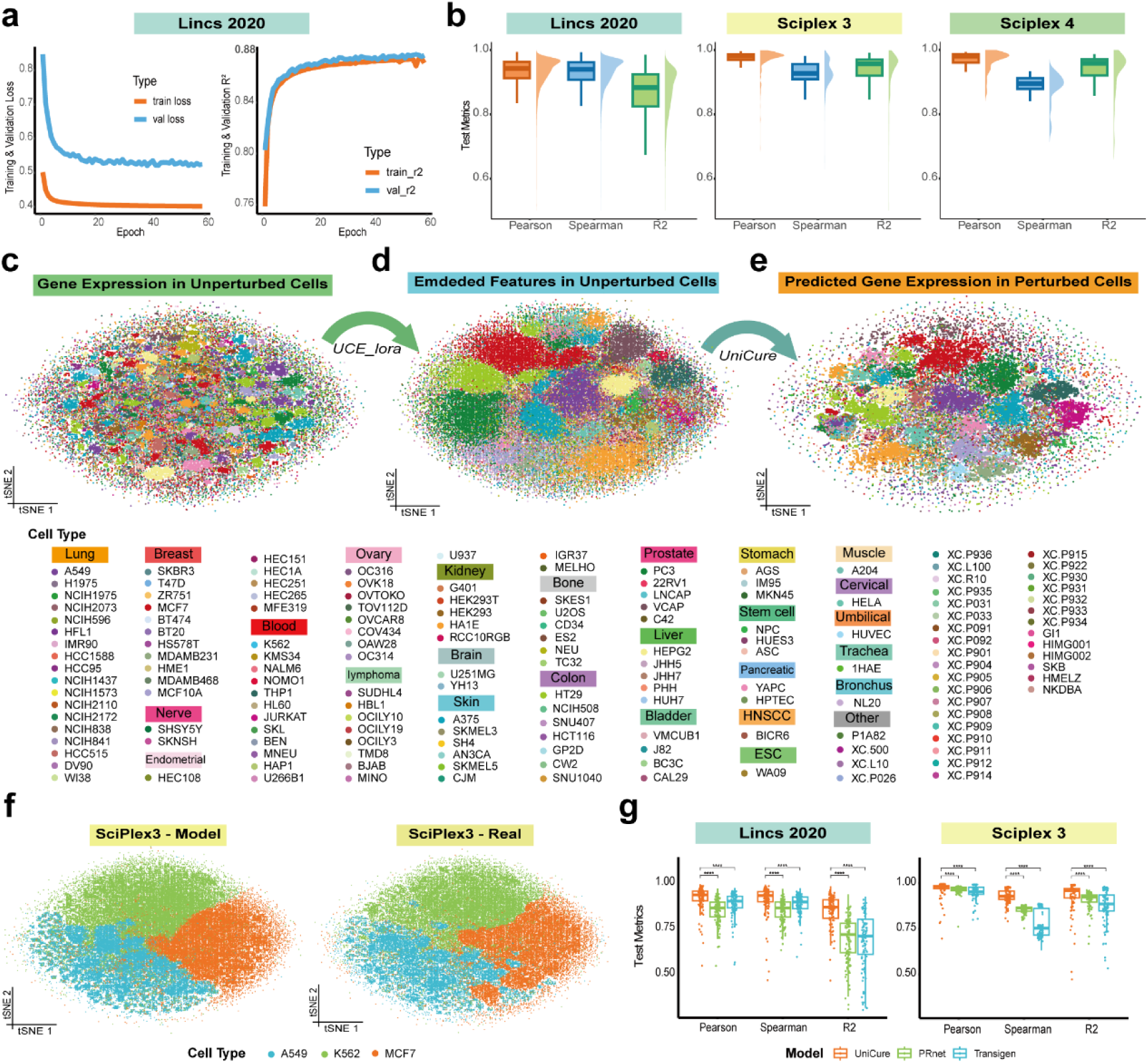
UniCure Accurately Predicts Drug-Induced Transcriptomic Perturbations at an individual level. **a**, Training and validation loss (left) and R² (right) across epochs during Stage 2 fine-tuning of UniCure on the LINCS2020 dataset. **b**, Performance of UniCure on held-out test sets of the LINCS2020, SciPlex3, and SciPlex4 datasets, evaluated using Pearson correlation, Spearman correlation, and R². t-SNE projections of LINCS2020 cells colored by cell type, showing unperturbed gene expression (**c**), embedded features after UCE_lora (**d**), and predicted perturbed gene expression by UniCure (**e**). **f**, t-SNE projections of perturbed cells in the SciPlex3 test set colored by cell type. Left, UniCure-predicted gene expression profiles. Right, experimentally observed profiles. **g**, Benchmark comparison of UniCure, PRnet, and TransiGen on the LINCS2020 and SciPlex3 test sets using Pearson correlation, Spearman correlation, and R². Statistical significance was assessed using the Wilcoxon rank-sum test. *P < 0.05; **P < 0.01; ***P < 0.001; ****P < 0.0001.

To gain mechanistic insights into UniCure’s performance, we visualized cellular representations in the model’s latent space using t-distributed Stochastic Neighbor Embedding (t-SNE). Prior to processing by the fine-tuned UCE module, cells from the same cell line (e.g., MCF7, red) frequently appeared fragmented into multiple distinct clusters, likely reflecting technical batch effects (Fig. 2c). In contrast, encoding through the LoRA-finetuned UCE component resulted in coherent clustering by cell line identity, indicating effective batch correction and the extraction of biologically meaningful features (Fig. 2d). We next examined the impact of predicted perturbations on these representations. Using representative unperturbed cells (n = 10 per cell line) and their matched test set drug–dose combinations, we found that most perturbed cells remained proximal to their original clusters, while some cell lines converged post-perturbation into shared regions—suggesting that UniCure captures convergent transcriptional programs induced by specific perturbations across diverse cell types. Conversely, other perturbed cells retained distinct separation, indicating preservation of cell-type–specific expression signatures under certain conditions (Fig. 2e).

Consistent with these findings, UniCure’s predicted transcriptomic profiles on the single-cell SciPlex3 dataset closely mirrored the real data in latent space. The t-SNE projections of predicted perturbations preserved the global cell-type structure and recapitulated the overall geometry observed in real perturbation profiles (Fig. 2f and Extended Data Fig. 2a-c), further supporting the model’s ability to learn biologically coherent and interpretable representations.

To benchmark UniCure against existing methods at the cell line level, we evaluated model performance on two public datasets: a randomly sampled subset of LINCS2020 (to mitigate computational burden), and the full SciPlex3 dataset. To ensure a fair comparison and avoid potential data leakage—given that prior models such as PRnet and Transigen may have been pretrained on these datasets—we retrained both baselines from scratch using the same data splits as UniCure. Using Pearson correlation, Spearman correlation, and R² as evaluation metrics, UniCure consistently outperformed both PRnet and TransiGen across datasets (Fig. 2g). Pairwise statistical comparisons using the Wilcoxon rank-sum test confirmed the significance of these improvements (****, P < 0.0001), further supporting UniCure’s enhanced ability to capture drug-induced transcriptomic perturbations. Notably, this analysis serves as an initial benchmark; a comprehensive evaluation against the original, publicly released versions of these models on primary PTC samples is presented in a subsequent section.

### UniCure Captures Drug Similarity, Dose-Responses, Mechanistic Signatures, and Combination Effects

To further interrogate UniCure’s ability to capture nuanced drug effects, we examined its predictions within specific cellular contexts. For example, visualizing predicted perturbations in the A375 melanoma cell line using t-SNE revealed that cells treated with the proteasome inhibitors Bortezomib and MG-132 clustered closely together, distinct from those treated with other compounds—reflecting their shared mechanism of action (MoA) (Fig. 3a). Notably, a similar clustering pattern was observed in the corresponding real perturbation data, supporting the model’s biological fidelity. We next examined dose–response relationships. At the highest tested concentration (20 µM), Bortezomib- and MG-132-treated cells formed a well-defined, concentration-specific cluster in both predicted and real expression spaces (Fig. 3b). In contrast, at the lowest dose tested (0.01 µM), UniCure predicted minimal transcriptional responses for most compounds, resulting in tightly grouped cell populations. However, this pattern was not as clearly observed in the real data, potentially reflecting platform-specific batch effects or reduced measurement sensitivity at low doses. Importantly, similar MoA- and dose-dependent clustering behaviors were recapitulated in additional cell lines, including MCF7 and PC3 (Extended Data Fig. 3a-d), further demonstrating the model’s capacity to capture compound- and concentration-specific transcriptomic signatures across diverse cellular backgrounds.

**Fig. 3:**
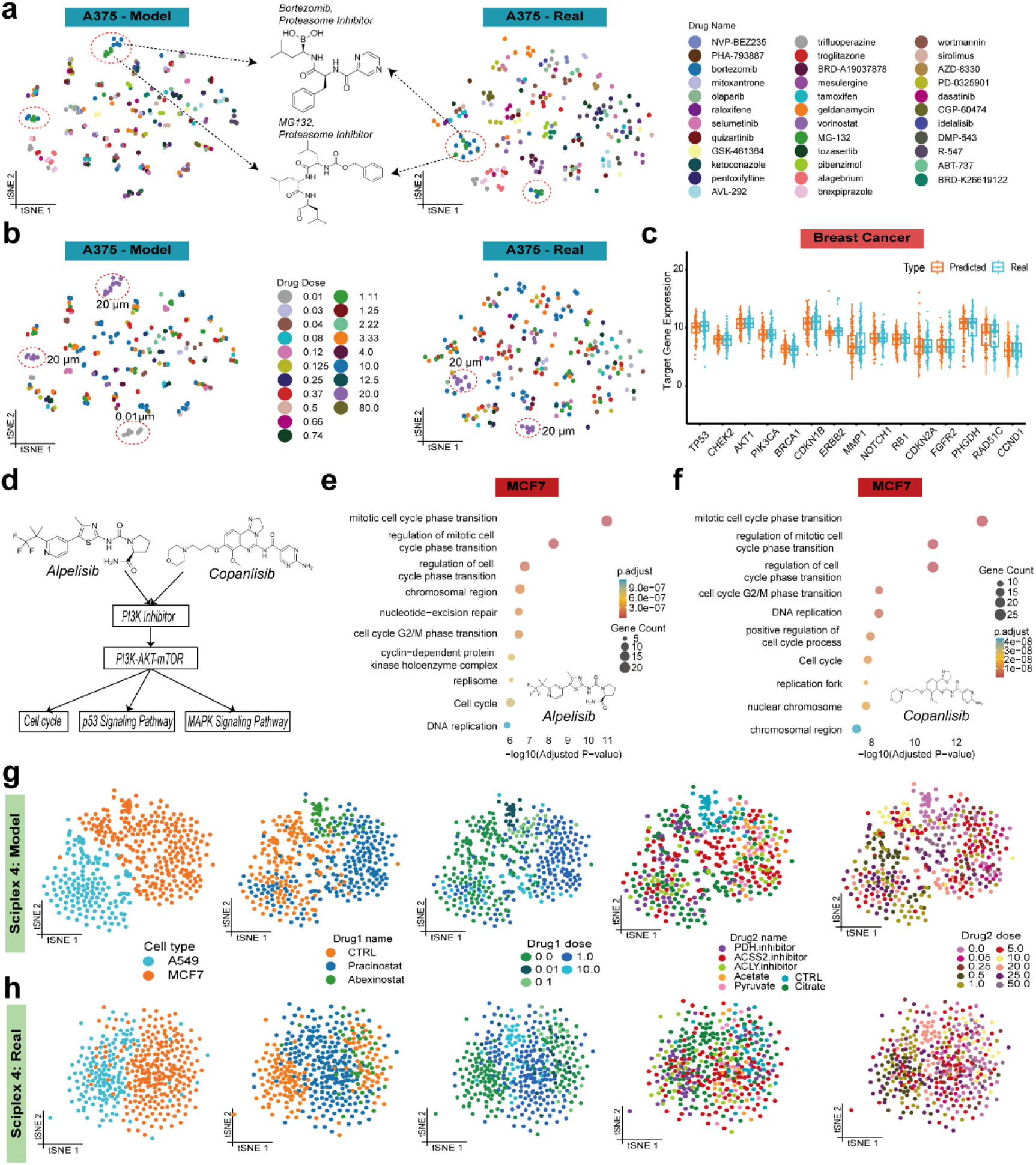
UniCure Captures Drug Similarity, Dose-Response, Mechanistic Signatures, and Combination Effects. **a**, UniCure-predicted (left) and experimentally observed (right) gene expression profiles in A375 cells, colored by compound identity. Bortezomib and MG132 (both proteasome inhibitors) are highlighted. **b**, Corresponding t-SNE projections colored by drug dose. 20 μM clusters for Bortezomib and MG132 are indicated in both model and real profiles. **c**, Comparison between UniCure-predicted and experimentally observed expression of breast cancer–relevant target genes across all breast cancer cell lines in the LINCS2020 dataset. **d**, Schematic representation of the shared mechanism of action for Alpelisib and Copanlisib. Both drugs inhibit PI3K, thereby affecting downstream PI3K–AKT–mTOR signaling and modulating the cell cycle, p53 signaling, and MAPK pathways. GO enrichment analysis of significantly downregulated genes in UniCure-predicted MCF7 profiles following treatment with Alpelisib (**e**) and Copanlisib (**f**). Top 10 enriched biological processes are shown, with dot size indicating gene count and color representing adjusted P value. t-SNE projections of UniCure-predicted (**g**) and experimentally observed (**h**) gene expression profiles in SciPlex4 dual-drug–perturbed cells, colored by cell type, first drug identity and dose, and second drug identity and dose (left to right).

We then assessed whether UniCure accurately predicts the modulation of cancer-relevant genes. We selected the top 15 most frequently reported target genes for breast cancer, lung adenocarcinoma, colon adenocarcinoma and prostate adenocarcinoma from the DisGeNET database^30^. Using LINCS2020 data, we examined the predicted expression distributions of these genes across all corresponding cancer-type cell lines following diverse drug perturbations, and compared them with experimentally observed expression profiles (Fig. 3c and Extended Data Fig4. a-c). UniCure’s predictions exhibited high concordance with the real data across cancer contexts, validating its capacity to model transcriptional responses of biologically meaningful targets across heterogeneous cellular backgrounds.

We extended this mechanistic analysis to the MCF7 cell line, which harbors a PIK3CA mutation. We predicted the effects of two PI3K inhibitors, alpelisib (FDA-approved for PIK3CA-mutant breast cancer) and copanlisib. Both drugs target the PI3K-AKT pathway, consequently affecting downstream signaling including the MAPK, cell cycle, and p53 pathways (Fig. 3d). UniCure predicted a predominantly suppressive effect on gene expression for both drugs in MCF7 cells (Extended Data Fig. 5a-b). Pathway enrichment analysis performed on significantly downregulated genes (adjusted p < 0.05, log2FoldChange < −0.3) revealed that pathways related to the cell cycle were among the most significantly enriched for both alpelisib and copanlisib (Fig. 3e, f). Furthermore, both MAPK and p53 signaling pathways were significantly enriched within the downregulated gene sets for both drugs (p.adj < 0.05, Supplementary Table 1, 2), consistent with their known downstream effects. Notably, this enrichment was not observed when applying the same drugs to the A375 melanoma cell line(p.adj < 0.05, Supplementary Table 3, 4), highlighting UniCure’s ability to capture cell-type–specific pathway responses. Notably, despite their distinct chemical structures (e.g., differing number of benzene rings), UniCure captured their similar functional impact, indicating that the model learns MoA based on biological response rather than simple structural similarity.

Finally, although large-scale public datasets for drug combinations remain limited, we leveraged the SciPlex4 dataset—which provides single-cell profiles for dual-compound perturbations—to explore UniCure’s potential in this domain. Visualizing UniCure’s predicted perturbation embeddings using t-SNE revealed clear separation and clustering of cells according to the identity of the first and second drugs in the combination (Fig. 3g). Similarly, distinct clusters emerged when coloring cells by the concentration of either compound, suggesting that UniCure can disentangle complex transcriptomic signatures arising from combinatorial treatments by capturing the individual and dose-specific effects of each component drug. Notably, similar clustering patterns were observed in the corresponding real perturbation profiles (Fig. 3h), further supporting the model’s ability to learn biologically coherent representations in multi-drug contexts.

### Establishment of PTC *in vitro* tumor models for bladder cancer and lung cancer

While large public cell line datasets allow models to learn foundational drug-cell interactions, their direct application to clinical scenarios is limited by the failure to recapitulate individual patient tumor biology and the tumor microenvironment (TME). To bridge this translational gap, we leveraged patient-derived tumor-like cell clusters (PTCs), which preserve crucial aspects of the original tumor, including immune components. A total of six fresh bladder (N=3) or lung (N=3) tumor samples were collected for PTC establishment. To maintain the integrity of tumor microenvironment components, we utilized the self-assembly of primary tumor cells with endogenous stromal and immune cells to establish three-dimensional clusters known as patient-derived tumor-like cell clusters in Matrigel-free conditions, as previously described^24^. PTCs typically achieved self-assembly within the first 2 to 3 days of culture and became denser around day 7 (Extended Data Fig. 6a). Hematoxylin and eosin (H&E) staining analysis revealed that bladder cancer PTCs exhibited solid spherical or nest-like structures, while lung cancer PTCs showed solid spherical or hollow ring-like structures, demonstrating high morphological similarities to surgically resected parental tumor samples (Extended Data Fig. 6b). The results of immunofluorescence staining indicated that PTCs broadly expressed pCK, a marker of epithelial cells, while the expression of CD45 and VIM suggested the presence of immune cells and fibroblasts or stromal cells, although the proportions varied significantly among different PTCs (Extended Data Fig. 6c). Notably, VIM-positive fibroblasts or stromal cells were typically located in the interior of the 3D structures of PTCs, in close interaction with pCK-positive epithelial cells. In contrast, CD45-positive immune cells were predominantly found on the surface of PTCs, with only Lu-3 samples exhibiting significant immune cell infiltration into the interior of PTCs, highlighting the variability in the tumor immune response characteristics of PTCs. These findings indicate that bladder and lung cancer PTCs can be rapidly constructed (within approximately 7 days) under our culture conditions, and that the PTCs are capable of maintaining the morphological characteristics of the source tissues and preserving the immune cell and stromal cell components of the tumor microenvironment. For each biological sample, a comprehensive 40-compound library—encompassing clinically approved chemotherapeutic and targeted therapeutic agents—was systematically screened. Meanwhile, compounds metadata and optimized concentrations are detailed in Supplementary Table S5. Transcriptomic profiling of drug-perturbed samples was subsequently performed to generate drug perturbation response signatures. Finally, we generated 123 PTC-derived drug perturbation data in LUAD and 123 PTC-derived drug perturbation data in BLCA. In addition, 79 triple-negative breast cancer (TNBC) PTC drug perturbation samples from Peking University People’s Hospital were included. These unpublished data, currently under independent investigation, were incorporated into UniCure training with institutional permission and informed patient consent. Together with 396 real-world patient-derived perturbation profiles—including samples from hematologic malignancies, breast cancers, and ovarian cancers—these datasets were used to fine-tune UniCure for personalized prediction.

### Efficient Fine-tuning on Patient-Derived Samples Enables Accurate Patient-Specific Prediction by UniCure

Recognizing the practical challenges of acquiring extensive patient perturbation data in clinical settings, we investigated how UniCure’s predictive performance scales with the amount of fine-tuning data for each cancer type. In the case of TNBC PTCs, fine-tuning UniCure on just 5% of the samples yielded a predicted versus actual gene expression correlation approaching 0.84; performance steadily improved with increasing training data, reaching nearly 0.9 with 80% of samples (Fig. 4a). Similar trends were observed in BLCA and LUAD PTCs: with only 5% of the data, UniCure achieved correlations of approximately 0.92 and 0.98, respectively, which further increased to values near 1.0 as more data were used (Fig. 4b,c).

**Fig. 4:**
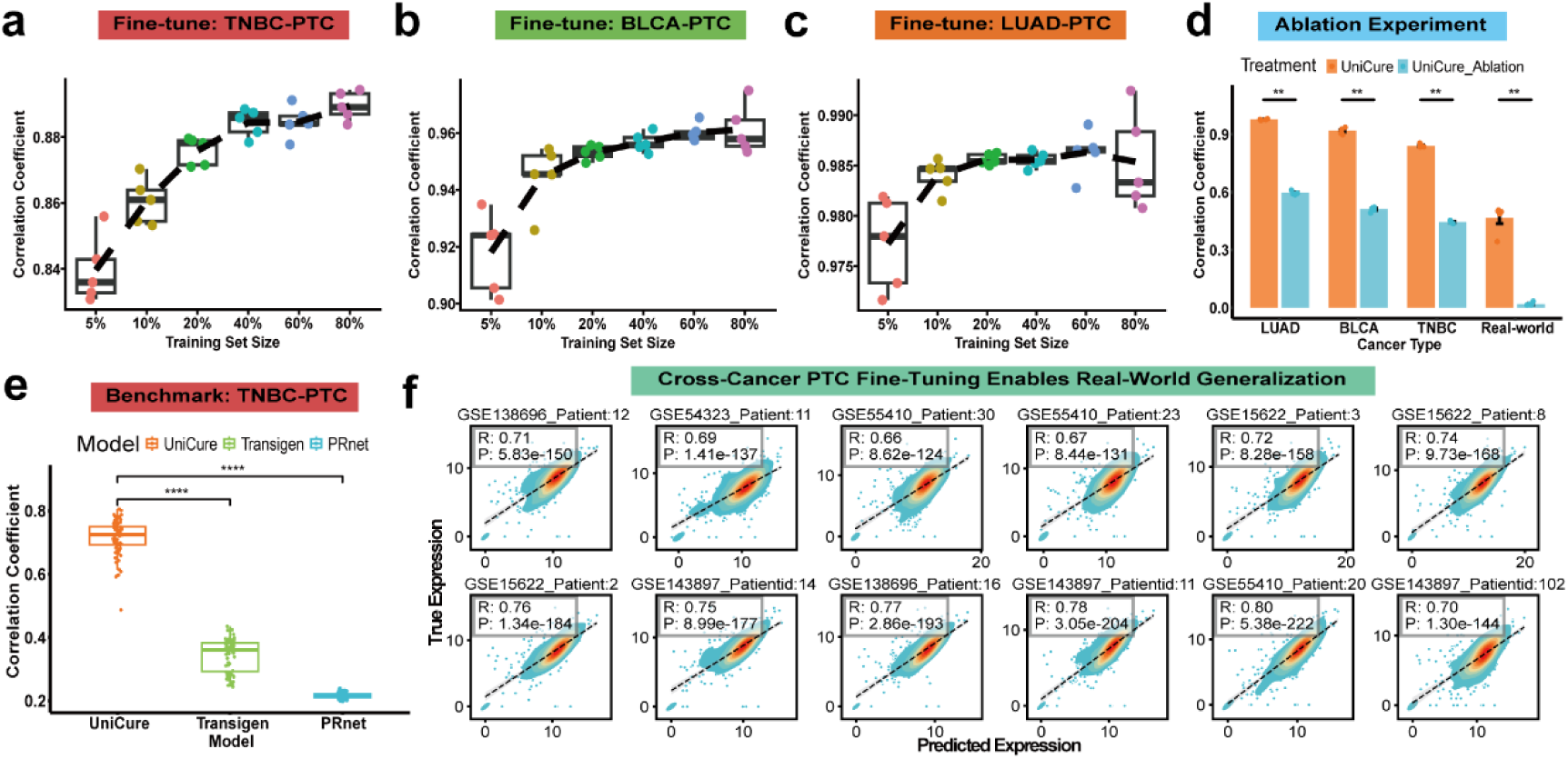
UniCure Enables Personalized Drug Response Prediction. **a–c**, UniCure was fine-tuned using increasing fractions of TNBC-PTC (**a**), BLCA-PTC (b), and LUAD-PTC (**c**) samples, and evaluated on held-out real-world data. Each point represents one of five replicates using different random seeds per training set size. Performance was assessed by correlation coefficient between predicted and observed expression. **d**, Ablation analysis comparing full UniCure and a version without pre-trained weights (UniCure_Ablation), each fine-tuned on 5% of data and evaluated on the remaining 95% across LUAD, BLCA, TNBC, and real-world datasets. Bars represent mean correlation (n = 5 seeds); error bars denote standard deviation; Wilcoxon rank-sum test, p < 0.01. **e**, t-SNE projections of perturbed cells in the SciPlex3 test set colored by cell type. Left, UniCure-predicted gene expression profiles. Right, experimentally observed profiles. **e**, Benchmark comparison of UniCure, TransiGen, and PRnet on TNBC-PTC test data. Wilcoxon rank-sum test, **** p < 0.0001. **f**, Scatter and density plots showing predicted versus true gene expression for 12 randomly selected real-world patient samples from hematologic malignancies, breast cancers, and ovarian cancers. UniCure was fine-tuned using TNBC-, BLCA-, and LUAD-PTC data.

To determine whether this strong performance critically depends on UniCure’s pre-training, we conducted an ablation study. We created an ablated version of UniCure by removing its pre-trained weights while keeping the architecture identical. This ablated model was trained from scratch using only 5% of PTC samples and evaluated on the remaining data. Across all three cancer types and real-world data, the ablated model yielded markedly lower correlations (ranging from 0.3 to 0.5) compared to the fully pre-trained UniCure, underscoring the essential role of foundational knowledge learned during pre-training in enabling accurate, data-efficient personalized predictions (Fig. 4c).

We then performed a rigorous comparison of UniCure against the officially released versions of TransiGen and PRnet on patient-level predictions. To prevent potential data leakage during evaluation and to ensure that UniCure could still learn patient-relevant transcriptional features, we adopted a cross-cancer fine-tuning strategy: when evaluating performance on one cancer type, UniCure was fine-tuned using PTC data from the other two types (e.g., fine-tuned on BLCA and TNBC PTCs when predicting LUAD). This strategy enabled the model to generalize across patient populations without relying on target-domain data during fine-tuning. In these cross-cancer experiments, UniCure consistently and significantly outperformed both TransiGen and PRnet across LUAD, BLCA, and TNBC PTC test sets. Pairwise statistical comparisons using the Wilcoxon rank-sum test confirmed the significance of UniCure’s improvements (****, adjusted P < 0.0001) (Fig. 4d and Extended Data Fig. 7a,b). Moreover, this cross-cancer strategy allowed us to assess UniCure’s ability to predict transcriptomic responses in clinically relevant, patient-derived samples across distinct cancer types (Extended Data Fig. 7c-e).

We further fine-tuned UniCure jointly on LUAD, BLCA, and TNBC PTC data and evaluated performance on the 396 real-world patient perturbation samples. The model achieved strong predictive performance, with average Pearson correlations around 0.7 across cancer types (Fig. 4e and Extended Data Fig. 7f). Notably, this performance was comparable to that of a baseline UniCure model fine-tuned using 80% of the real-world patient data—highlighting UniCure’s ability to generalize from curated datasets to external patient cohorts with high data efficiency(Extended Data Fig. 7g).

### UniCure-derived Personalized Drug Rankings Stratify Patients and Reflect Underlying Tumor Biology

To further evaluate UniCure’s potential for clinical translation, we assessed its ability to generate personalized drug response predictions using patient data from the UCSC Xena Browser^31^. We focused on eight cancer types where paired tumor and adjacent normal tissue transcriptomic data were available(Method). For each patient’s tumor sample, we employed UniCure to predict the transcriptomic perturbations induced by nearly 5,000 of common FDA-approved drugs, clinical trial drugs, and preclinical tool compounds.

To quantify the potential therapeutic effect of each drug for a given patient, we calculated the Pearson correlation between the predicted drug-induced gene expression changes and the patient’s intrinsic tumor-versus-normal tissue gene expression differences. A lower (more negative) correlation coefficient signifies a greater predicted ability of the drug to reverse the tumor-associated transcriptomic signature (Method). Based on these correlation scores, we ranked about 5,000 compounds for each patient, generating a high-dimensional personalized drug ranking vector representing predicted therapeutic priorities (Fig. 5a).

**Fig. 5.**
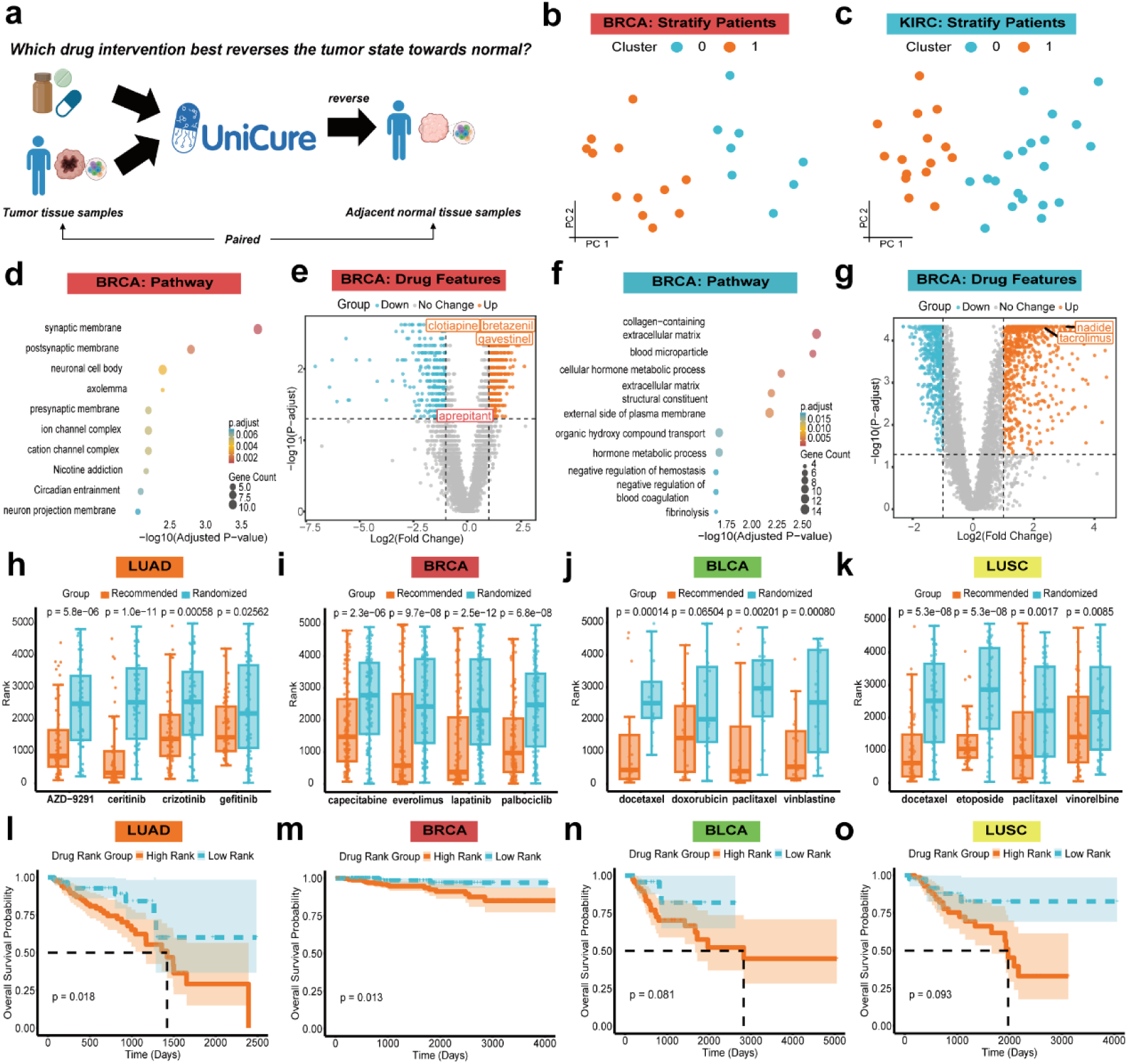
Personalized Drug Rankings by UniCure Stratify Patients, Reflect Tumor Biology, and Correlate with Clinical Outcomes. **a**, Schematic of UniCure’s personalized drug prioritization strategy based on selecting compounds that most effectively reverse tumor transcriptomic profiles toward matched adjacent normal tissue. **d–g**, Each patient was represented by the log_10_-normalized ranking scores of ∼5,000 small-molecule compounds predicted by UniCure. K-means clustering was applied to the resulting feature matrix for advanced BRCA (**b**) and KIRC (c) cohorts. GO pathway enrichment (**d, f**) and differential drug ranking (**e, g**) between UniCure-stratified patient clusters in BRCA (**d–e**) and KIRC (**f–g**), based on differentially expressed genes and drug prioritization profiles, respectively. **h–k**, Predicted rank distributions of clinically relevant therapies across public cohorts of LUAD (**h**), BRCA (**i**), BLCA (**j**), and LUSC (**k**) patients, compared to randomized controls; significance assessed by Wilcoxon rank-sum test. **i–o**, Kaplan–Meier survival analysis for LUAD (**i**), BRCA (**m**), BLCA (**n**), and LUSC (**o**) patients, stratified by whether the administered drug ranked in the top 30% (High Rank) or bottom 70% (Low Rank) of UniCure’s predicted rankings.

We hypothesized that these personalized drug ranking vectors could capture latent biological heterogeneity and serve as a basis for novel patient stratification. We focused this analysis on patients with late-stage (Stage III/IV) disease, where therapeutic decisions are most critical, selecting breast cancer (BRCA, n=36) and clear cell renal cell carcinoma (KIRC, n=20) cohorts with sufficient sample sizes (n>20). Applying K-means clustering to the personalized drug ranking vectors for each cancer type, both the elbow method and silhouette score analysis consistently indicated an optimal cluster number of two (k=2) for both BRCA and KIRC (Fig. 5b, c and Extended Data Fig. 8a, b).

We then investigated the biological basis underlying these data-driven patient strata. In BRCA, pathway enrichment analysis of DEGs between the two clusters revealed a striking overrepresentation of pathways related to neuronal processes, including ‘synaptic membrane’ and ‘postsynaptic membrane’ (Fig. 5d Supplementary Table S6). Consistent with this finding, analysis of the drugs exhibiting the most significantly different rankings between the two BRCA clusters (ranked by adjusted p-value) identified compounds predominantly associated with modulating neuronal systems. These included agents targeting the neurotransmitter systems (e.g., clotiapine, bretazenil, fenthion, gavestinel, WB-4101), endocannabinoid system (e.g., mead-ethanolamide, AM-404, BML-190), and neuropeptide/neuromodulator receptors (e.g., talnetant, antalarmin) (Fig. 5e and Supplementary Table S7). This observation resonates strongly with recent findings, including a study describing a crucial neuro-cancer interaction axis in breast cancer metastasis involving sensory neuron activation, substance P (SP) release, and subsequent TLR7-mediated pro-metastatic reprogramming in adjacent cancer cells via TACR1 signaling^27^. Importantly, this axis was shown to be pharmacologically targetable, for instance, by the TACR1 antagonist Aprepitant. Examining Aprepitant within our framework, we found its ranking difference between the two BRCA patient clusters showed a strong trend towards significance (Fig. 5e, raw p = 0.0124, adjusted p = 0.0545). This borderline significance, despite multiple hypothesis correction across ∼5000 drugs, suggests that UniCure’s personalized rankings may indeed capture differential sensitivity related to this neuro-cancer axis, potentially identifying a patient subgroup where targeting TACR1 could be particularly relevant.

Turning to the KIRC cohort, pathway enrichment analysis of genes distinguishing the two patient clusters revealed three dominant biological themes: extracellular matrix (ECM)/tumor microenvironment interactions, blood coagulation and fibrinolysis systems, and diverse metabolic processes (Fig. 5f, and Supplementary Table S8). These findings are highly consistent with the known biology of KIRC, a vascularized tumor subtype characterized by extensive metabolic reprogramming and ECM remodeling^32^. Analysis of differentially ranked drugs between the two clusters further corroborated these biological differences (Fig. 5g). Tacrolimus, an immunosuppressant known to enhance mTORC1/2 signaling, exhibited significantly divergent rankings; notably, prior studies suggest it may promote RCC initiation and progression in renal transplant recipients^33^. Nadide (NAD⁺), a central metabolic coenzyme, was also differentially ranked, reflecting the metabolic divergence between clusters. Additional compounds further emphasized these biological themes, including Pirfenidone (anti-fibrotic, ECM modulation), Ticlopidine (anti-platelet, involved in hemostasis), and several agents targeting metabolic and redox pathways such as Bromebric acid (lipid metabolism), Coumaric acid and Eugenitol (oxidative stress balance), PPT (hormonal metabolism), and Indoximod (amino acid metabolism)) (Supplementary Table S9). The alignment between drug mechanisms and the biological signatures separating patient clusters highlights UniCure’s capacity to generate biologically meaningful drug rankings tailored to the functional landscape of distinct patient subgroups.

### Clinical Validation of UniCure’s Personalized Drug Recommendations via Known Therapies and Patient Survival

Having established that UniCure-derived rankings can stratify patients biologically, we next sought to validate whether the rankings of specific, clinically relevant drugs align with known therapeutic strategies and patient outcomes. We first examined the rank distribution of well-established targeted therapies within their indicated cancer types among the personalized rankings generated for paired tumor-normal samples. In LUAD, for instance, Osimertinib (AZD-9291), a potent EGFR tyrosine kinase inhibitor (TKI) frequently used in EGFR-mutant lung cancer, ranked within the top 1,000 predicted therapies on average across patients. To assess statistical significance, we compared the observed rank distribution of Osimertinib against a null distribution generated by randomly permuting each patient’s drug ranking vector. This revealed a highly significant enrichment of Osimertinib towards higher-priority ranks (one-sided Wilcoxon rank-sum test, p = 5.8 x 10⁻⁶). Similar significant enrichment towards higher-priority rankings was observed for other established LUAD therapies, including the ALK inhibitor Ceritinib, the ALK/ROS1/MET inhibitor Crizotinib, and the EGFR TKI Gefitinib (Fig. 5h).

We extended this analysis across multiple cancer types, evaluating the rankings of specific targeted agents relevant to each malignancy (e.g., pertinent TKIs, endocrine therapies). Significant enrichment of appropriate targeted therapies among the higher-priority UniCure rankings was confirmed in BRCA, LUSC, BLCA, KIRC, and LIHC (Fig. 5i-k, Extended Data Fig. 8c-f). However, this enrichment was not statistically significant for the drugs examined in COAD and PRAD. This could potentially be attributed to several factors, including the possibility that the dominant biological drivers or drug sensitivities in these specific cohorts are less well represented within the foundational cell line and single-cell data used for UniCure’s pre-training, or limitations in capturing the full complexity of these diseases and their treatment responses solely through baseline transcriptomics.

To directly link UniCure’s predictions to clinical benefit, we investigated the association between the predicted ranking of a patient’s actually administered therapy and their survival outcome. Due to the sparsity of treatment records combined with paired normal samples, we expanded our cohort to include all patients with available treatment information, using the average expression profile of adjacent normal tissues within that cancer type as the reference for calculating the drug ranking score (as described previously). Given the very low number of PRAD patients with usable treatment data (n=8), this cohort was excluded from the survival analysis. For each remaining cancer type, patients were stratified into two groups based on the UniCure-predicted rank percentile of their documented first-line therapy: a “High-Priority Treatment” group (drug ranked within the top 30th percentile) and a “Low-Priority Treatment” group (drug ranked below the 30th percentile).

Kaplan-Meier survival analysis revealed significantly improved overall survival for patients in the High-Priority Treatment group compared to the Low-Priority Treatment group in LUAD (log-rank p = 0.0038) and BRCA (log-rank p = 0.0088) (Fig. 5l, m). Trends towards improved survival were also observed in BLCA (p = 0.081) and LUSC (p = 0.093) (Fig. 5n, o). Conversely, no significant survival difference between the groups was observed in COAD, KIRC, and LIHC (Extended Data Fig. 8g-i). Potential reasons for this lack of significance in certain cohorts may include the necessary approximation using average normal tissue profiles, limited sample sizes with complete treatment and follow-up data, heterogeneity within the recorded treatments, the influence of unrecorded subsequent therapies, or that the primary drivers of outcome in these specific patient groups are less correlated with the transcriptomic response signature captured by UniCure for the administered drugs.

Illustrative examples from the LUAD cohort with paired tumor-normal samples underscore the potential clinical relevance of UniCure’s findings (Table 1). For instance, patient 49-6743 (female, Stage IIIA, 81y), who received Pemetrexed—a standard chemotherapy agent—and survived 1621 days, had the drug ranked in the top ∼10% by UniCure, placing her in the High-Priority Treatment group. In contrast, patient 50-6595 (female, Stage IIIA, 74y), who received Carboplatin and survived only 189 days, had the drug ranked in the bottom ∼1% by UniCure, corresponding to the Low-Priority Treatment group. Similar trends were observed in BRCA, BLCA, and LUSC cohorts (Supplementary Table S10-12), further supporting the alignment of high-priority predictions with better patient outcomes. However, this relationship was less evident in COAD, KIRC, and LIHC cohorts (Supplementary Table S13-15), likely due to cohort-specific heterogeneity, gaps in training data, and limitations in treatment annotations. These findings demonstrate the potential of UniCure’s framework while highlighting the need for further optimization to improve its applicability across diverse cancer types.

**Table 1.** Clinical characteristics and UniCure drug recommendations for LUAD patients. Clinical and survival information for matched LUAD patients from TCGA, along with actual treatment and UniCure-based drug recommendation status (✔: top 30% prioritized drug; 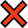: not recommended).

| PATIENT ID | CANCER | PATHOLOGIC STAGE | GENDER | DAYS TO BIRTH | DEATH? | DAYS TO LAST FOLLOW UP | DRUG NAME | DRUG RANK(%) | UNICURE RECOMMENDATION (TOP 30%) |
| --- | --- | --- | --- | --- | --- | --- | --- | --- | --- |
| TCGA.49.6743 | LUAD | Stage IIIA | FEMALE | 81 | 0 | 1621 | pemetrexed | 10 | ✓ |
| TCGA.44.2665 | LUAD | Stage IIB | FEMALE | 55 | 0 | 1301 | pemetrexed | 7.5 | ✓ |
| TCGA.44.2662 | LUAD | Stage IB | MALE | 65 | 0 | 1280 | paclitaxel | 97.3 | ✗ |
| TCGA.44.3396 | LUAD | Stage IIIA | FEMALE | 74 | 0 | 1130 | pemetrexed | 23.1 | ✓ |
| TCGA.44.6146 | LUAD | Stage IIB | MALE | 64 | 0 | 728 | pemetrexed | 19.1 | ✓ |
| TCGA.49.6742 | LUAD | Stage IIA | MALE | 70 | 0 | 445 | pemetrexed | 7.0 | ✓ |
| TCGA.73.4676 | LUAD | Stage IIA | MALE | 45 | 0 | 164 | cisplatin | 19.7 | ✓ |
| TCGA.50.6595 | LUAD | Stage IIIA | FEMALE | 74 | 1 | 189 | carboplatin | 99.6 | ✗ |
| TCGA.55.6979 | LUAD | Stage IIB | FEMALE | 59 | 1 | 237 | paclitaxel | 56.7 | ✗ |
| TCGA.50.5936 | LUAD | Stage IIIA | MALE | 58 | 1 | 257 | paclitaxel | 63.7 | ✗ |
| TCGA.50.5930 | LUAD | Stage IIIA | MALE | 47 | 1 | 282 | docetaxel | 8.1 | ✓ |
| TCGA.49.4490 | LUAD | Stage IIIA | FEMALE | 45 | 1 | 385 | cisplatin | 99.9 | ✗ |
| TCGA.55.6970 | LUAD | Stage IIIA | FEMALE | 67 | 1 | 464 | cisplatin | 56.8 | ✗ |
| TCGA.55.6982 | LUAD | Stage IIB | FEMALE |  | 1 | 995 | vinorelbine | 87.3 | ✗ |
| TCGA.38.4632 | LUAD | Stage IV | MALE | 42 | 1 | 1357 | carboplatin | 96.8 | ✗ |
| TCGA.50.5933 | LUAD | Stage IIIB | MALE | 72 | 1 | 2393 | paclitaxel | 72.1 | ✗ |

### UniCure identified active natural products against human cancer

Natural products (NPs) are prized for their chemical diversity, therapeutic potential, and favorable safety profiles, playing pivotal roles in precision oncology^34^. Nearly half of existing anticancer medications originate either directly or through modification from NPs^35^. To further prove the value of UniCure, we predicted the transcriptomic perturbations profile induced by the natural compounds available in our laboratory (2019 compounds) to identify potential novel compound candidates against human cancer. The profile-based approach identifies potential therapeutics by selecting compounds that induce gene expression patterns opposing disease-associated signatures (Fig. 6a). This method operates on the principle that drugs showing negative correlation between their perturbation signatures and disease-specific gene expression profiles are likely to exhibit therapeutic efficacy against the condition^36^. Here, we *in silico* screened the natural compound library available in our laboratory to identify potential compound candidates across three cancer type perturbing 3 cell lines, including triple-negative breast cancer (MDA-MB-231), lung adenocarcinoma (A549), and bladder cancer (5637). Note that these compounds were not included in the training set. We created disease gene expression signatures across triple-negative breast cancer (TNBC), lung adenocarcinoma (LUAD) and bladder cancer (BLCA) by comparing RNA-sequencing (RNA-Seq) gene expression from tumors and adjacent normal tissues, using data downloaded from TCGA (except for TNBC, which the expression data downloaded from FUSCC-TNBC cohort). After subsetting to the set of LINCS landmark genes, the signatures of TNBC, LUAD and BLCA included 169, 143 and 159 differentially expressed (DE) genes in tumors compared to the normal tissues of each (Supplementary table S16-S18), respectively and these signatures could accurately classify tumors and adjacent normal tissues (Extended Data Fig. 9).

**Fig. 6:**
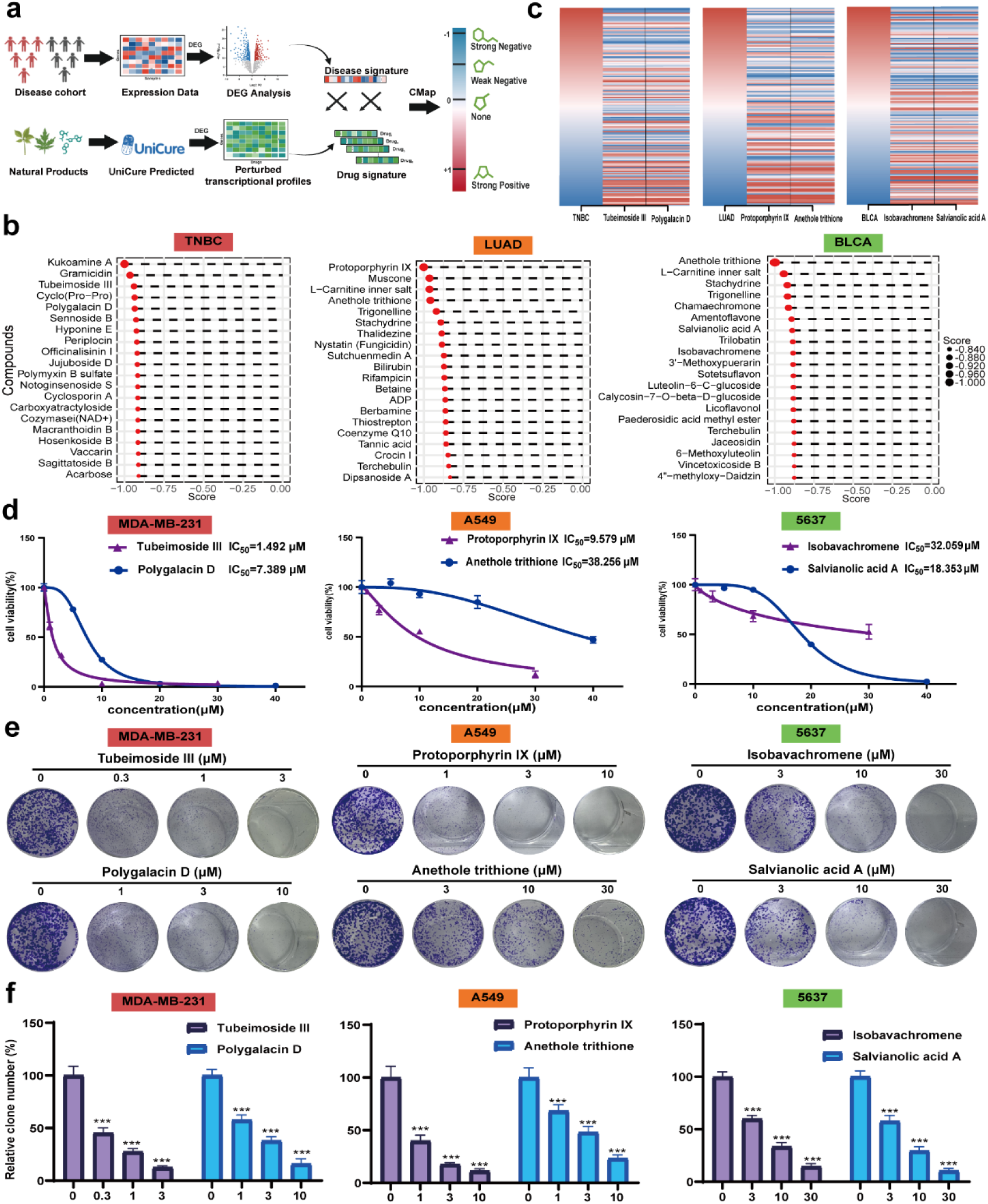
Experimental Validation of UniCure-Predicted Therapeutic Candidates. (**a**) Overview of the UinCure-based drug identification (**b**) Top 20 compounds for each cancer based on the connectivity score. (**c**) Reversal relationship between disease signature (TNBC/LUAD/BLCA) and drug candidate signatures. (**d**) Drug efficacy in three cancer lines, including MDA-MB-231, A549 and 5637, measured by cell proliferation assays after 48 h treatment. (**e-f**) Drug candidates demonstrated inhibition of colony formation in MDA-MB-231 cells, A549 and 5637 cells. Data represents mean ± standard error of the mean (SEM) from three independent biological replicates. ∗P < 0.05; ∗∗P < 0.01; ∗∗∗P < 0.001.

These signatures were compared against individual predicted drug gene expression profile (TNBC: Predicted drug gene expression profiles in MDA-MB-231 cell; LUAD: Predicted drug gene expression profiles in A549 cell; BLCA: Predicted drug gene expression profiles in 5637 cell), resulting in one connectivity score for each compound profile for each cancer (scores range from −1 to 1, Supplementary table S19-S21). Interestingly, many of the top-ranked compounds exhibit biological activity against TNBC, LUAD, and BLCA and top 20 compounds were shown in Fig. 6b. Two natural compounds at the top rank (rank ≤ 10) for each cancer were chosen as the candidate set for validation (Tubeimoside III and Polygalacin D for TNBC; Protoporphyrin IX and Anethole trithione for LUAD; Isobavachromene and Salvianolic acid A for BLCA). We evaluated the anti-cancer activity of candidate compounds against three cell lines utilizing CCK-8 assays. We observed that all six compounds significantly inhibited cancer cells viability in a concentration-dependent manner. The IC_50_ values were calculated to determine the effectiveness of the compounds against the specific cancer cell lines. In MDA-MB-231 cells, Tubeimoside III and Polygalacin D demonstrated potent cytotoxic effects, with IC_50_ values of 1.492 µM and 7.389 µM, respectively; Protoporphyrin IX and Anethole trithione exhibited IC_50_ values of 9.579 µM and 38.256 µM, respectively, in A549 cells; Isobavachromene and Salvianolic acid A exhibited IC_50_ values of 32.059 µM and 18.353 µM, respectively, in 5637 cells (Fig. 6c). As shown in Fig. 6d, these compounds also demonstrated obvious inhibitory effects on the corresponding tumor cells. Moreover, the clonogenic assay also revealed a concentration-dependent inhibition of colony formation by the two compounds in each individual cancer cells. At 1 µM, Tubeimoside III and Polygalacin D notably decreased colony formation abilities in MDB-MA-231 cells by 27.38% (P < 0.0001) and 57.59% (P < 0.0001) (Fig. 6e), respectively. Additionally, the suppressive effect of Protoporphyrin IX and Anethole trithione on the colony formation of A549 cells exhibited a clear concentration dependency. At concentrations of 1 µM and 3 µM, the inhibition rates of these two compounds were 39.64% (P < 0.0001), 17.44% (P < 0.0001) and 68.21% (P < 0.0001), 48.00% (P < 0.0001) for A549 cells. Moreover, in 5637 cells, the Isobavachromene and Salvianolic acid A exhibited decreasing reductions of 59.84% (P < 0.0001), 33.37% (P < 0.0001) and 57.63% (P < 0.0001), 29.42% (P < 0.0001) respectively at concentrations of 3 µM and 10 µM (Fig. 6f). Hence, the results above emphasize the efficacy of these candidates in inhibiting the growth of particular cancer cell lines, indicating their promising therapeutic potential in cancer treatment.

## Discussion

Predicting personalized drug responses from transcriptomic data represents a critical challenge in advancing precision oncology. Existing computational models often struggle to translate findings from the simplified cell line systems to the complicated patient tumors. Here, we introduce UniCure, the first pre-trained foundation model integrating both omics and chemical foundation models (UCE and Uni-mol) to predict how drugs alter transcriptomes across diverse biological contexts, facilitating personalized cancer therapy and drug prioritization. Our comprehensive evaluations demonstrate that UniCure not only achieves state-of-the-art performance in predicting transcriptomic perturbations across diverse datasets but also generates clinically relevant insights with potential translational value.

A key strength of UniCure lies in its architecture, leveraging powerful pre-trained foundation models for both drug (Uni-mol) and cell (UCE) representations. The incorporation of LoRA within the UCE module facilitated efficient adaptation of the cellular encoder, preserving foundational biological knowledge while allowing flexibility. Subsequently, the framework was further adapted for personalization through fine-tuning on patient-derived data (PTCs and real-world samples), adding specific trainable parameters to capture individual tumor characteristics. This approach proved highly effective, enabling rapid adaptation to new patient contexts using remarkably small datasets (as few as 10 PTC samples). While the high predictive correlations observed may partly reflect inherent similarity among samples within each cancer type, our ablation studies confirmed that UniCure’s performance is not solely attributable to dataset redundancy. The observed data efficiency critically depends on the pre-trained weights, underscoring the importance of leveraging foundational knowledge for personalized prediction in settings where patient-specific data are limited. Furthermore, the novel FlexPert module adeptly handled flexible drug inputs, and the use of MMD loss successfully addressed the challenge of unpaired perturbation data, enhancing robustness and generalization compared to methods reliant solely on paired samples or simpler loss functions. These technical innovations collectively contribute to UniCure’s superior performance over existing methods like TransiGen and PRnet on both cell line benchmarks and, more importantly, on patient-derived PTC and real-world perturbation data.

Beyond predictive accuracy, UniCure demonstrated a capacity to capture nuanced biological information. Its ability to discern drugs with similar mechanisms of action (e.g., proteasome inhibitors), resolve dose-dependent effects, and predict expression changes in key cancer-associated genes suggests that the model learns biologically meaningful relationships rather than relying on superficial correlations. Notably, UniCure accurately captured the similar functional consequences of structurally distinct compounds targeting the same pathway, such as the PI3K inhibitors alpelisib and copanlisib. The model also effectively disentangled transcriptomic responses to combinatorial drug perturbations. Furthermore, key UniCure predictions were validated not only against public cohort data but also experimentally *in vitro*, reinforcing its role as a biologically grounded and translationally relevant framework.

The application of UniCure to the public clinical cohorts yielded particularly compelling results regarding its clinical potential. The framework’s ability to generate personalized drug rankings based on reversing tumor transcriptomic signatures allowed for data-driven stratification of late-stage BRCA and KIRC patients into distinct subgroups. Importantly, the biological pathways and differentially ranked drugs distinguishing these clusters aligned remarkably well with known tumor biology (neuronal signaling in BRCA, ECM/metabolism/coagulation in KIRC), suggesting these *in silico* rankings capture underlying functional heterogeneity. This approach could potentially uncover novel patient subtypes or biomarkers predictive of response to specific therapeutic classes.

Furthermore, the validation against known clinical practices provides strong evidence for UniCure’s relevance. The preferential high ranking of established targeted therapies (e.g., Osimertinib in LUAD) within their indicated cancer types suggests the model has learned clinically meaningful drug-disease associations. Most significantly, the correlation between the UniCure ranking of a patient’s actual administered therapy and their overall survival in LUAD and BRCA offers a direct link between the model’s predictions and patient outcomes. While this association requires further validation, it strongly suggests that UniCure captures information pertinent to treatment efficacy and could potentially inform therapeutic decision-making.

Despite these promising results, several limitations should be acknowledged. UniCure primarily relies on transcriptomic data, neglecting other crucial biological layers like genomics, proteomics, and epigenetics that influence drug response. While PTCs offer a significant improvement over cell lines, they remain an *ex vivo* system and may not fully recapitulate the complex TME dynamics or long-term treatment effects *in vivo*. Another current limitation is the relative scarcity of large-scale, high-quality public datasets specifically profiling drug combinations, particularly in patient-derived models or clinical settings. Although UniCure’s architecture, featuring the FlexPert module, is explicitly designed to process multiple drug inputs, and initial results on the SciPlex4 dual-perturbation data are encouraging, further validation and training on more extensive combination datasets are needed. As such data becomes increasingly available from preclinical screens or clinical trials employing combination therapies, UniCure is well-positioned to leverage this information to predict synergistic or antagonistic effects, a critical step towards designing more effective combination strategies. The public clinical cohorts survival analysis, while encouraging, relied on retrospective data with inherent heterogeneity in treatments and relied on average normal profiles for many samples, potentially confounding the results.

Future work should focus on integrating multi-omics data to provide a more holistic view of the cell state and drug response. Enhancing the model’s ability to explicitly represent and predict interactions with specific TME components is another critical direction. Expanding the drug library and validating the framework’s ability to predict responses to novel chemical entities not seen during pre-training would further strengthen its utility. Ultimately, prospective clinical studies are necessary to rigorously evaluate whether UniCure-guided therapy selection can improve patient outcomes compared to standard-of-care approaches.

In conclusion, UniCure represents a significant advancement in predicting personalized drug responses by effectively integrating large-scale foundation models with patient-specific data and PEFT strategies. Its demonstrated accuracy, biological insight, data efficiency, and correlation with clinical outcomes highlight its potential as a powerful tool to accelerate drug discovery, optimize clinical trial design through predictive stratification, and ultimately guide personalized therapeutic strategies in oncology.

## Method

### The L1000 dataset

The LINCS consortium provides high-throughput gene expression profiles under diverse pharmacological perturbations. In this study, we leveraged the L1000 assay, which quantifies 978 landmark genes. Our analysis focused on the CMAP LINCS 2020 release, specifically Level 3 data (processed perturbation profiles alongside matched controls). To ensure consistency, we applied stringent filters: retained only 24-hour exposure experiments across all valid concentration ranges (excluding ambiguous or missing dose annotations); removed compounds with invalid SMILES strings (non-parsable by RDKit); each perturbed profile was paired with its corresponding unperturbed control and unmatched entries were discarded. The final curated dataset comprised 501449 perturbation-control pairs, spanning 22593 compounds and 166 cell lines, providing a comprehensive resource for downstream analysis.

### The sci-Plex3 and sci-Plex4 dataset

We utilized two complementary sci-Plex series single-cell transcriptomic datasets downloaded from PerturBase database the for model evaluation^37^. The sci-Plex3 dataset (original release) captured transcriptional responses of three human cancer cell lines (MCF7, K562, and A549) to 188 individual compounds through high-throughput screening. For broader validation, we incorporated the updated sci-Plex4 dataset, which extends this paradigm by profiling responses to drug combinations in A549 and MCF7 cell lines. All datasets underwent standardized quality control using Seurat with the following parameters: minimum gene detection threshold: ≥400 genes per cell; maximum mitochondrial content: ≤20%; top 1,000 highly variable genes using FindVariableFeatures function; exclusion of compounds with RDKit-unparsable SMILES strings; strict matching of perturbed samples with corresponding controls. The final curated datasets comprised: sci-Plex3: 2255 perturbation-control pairs (3 cell lines; 188 compounds); sci-Plex4: 204 perturbation-control pairs (2 cell lines; 9 compounds).

### Real-world patient drug perturbation profiles

The real-world patient drug perturbation profiles (transcriptomic profiles of the same patient sample pre- and post-drug administration) were collected from the Cancer Treatment Response gene signature DataBase (CTR-DB 2.0)^38^. The CTR-DB project is the first data resource for clinical transcriptomes with cancer treatment response, and meanwhile supports various data analysis functions. Our curation pipeline enforced strict paired-sample requirements (same patient, pre/post treatment), yielding 396 (N=198 patients) high-confidence patient perturbation profiles. The complete dataset composition is cataloged in Supplementary table S22.

### Processing of Public Clinical Cohorts

Public clinical cohort data were obtained from the UCSC Xena Browser TCGA Hub, covering eight cancer types: lung adenocarcinoma (LUAD), breast invasive carcinoma (BRCA), bladder urothelial carcinoma (BLCA), lung squamous cell carcinoma (LUSC), kidney renal clear cell carcinoma (KIRC), colon adenocarcinoma (COAD), prostate adenocarcinoma (PRAD), and liver hepatocellular carcinoma (LIHC). For initial analysis, we specifically selected patients with paired tumor and adjacent normal tissue transcriptomic profiles to support the modeling of individual drug response, including BLCA (n=19), BRCA (n=112), COAD (n=41), KIRC (n=72), LIHC (n=50), LUAD (n=57), LUSC (n=49), and PRAD (n=52).

For survival analyses, the dataset was expanded to include patients without matched normal tissue but who had complete follow-up information, overall survival data, and treatment records, resulting in a cohort of BLCA (n=102), BRCA (n=533), COAD (n=97), KIRC (n=59), LIHC (n=31), LUAD (n=165), and LUSC (n=128). This ensured a comprehensive and clinically meaningful evaluation of model-predicted treatment efficacy across diverse real-world scenarios.

### Disease signatures

Bladder cancer and lung adenocarcinoma cohorts were obtained from The Cancer Genome Atlas (TCGA), containing RNA-seq data from tumor and adjacent normal tissues. For triple-negative breast cancer (TNBC), we analyzed 360 TNBC patients (360 tumor tissues / 88 normal tissues) with RNA-seq data from a 465-patient cohort established at Fudan University Shanghai Cancer Center (FUSCC)^39^. Disease signatures were derived using limma package with default parameters by comparing tumor vs. matched normals, selecting genes with |log2FC| > 1 and P < 0.05. Signatures were refined by intersecting with LINCS landmark genes for subsequent compound screening. A total of 133 up-regulated genes and 36 down-regulated genes for TNBC were identified; 77 up-regulated genes and 76 down-regulated genes for LUAD were identified; and 95 up-regulated genes and 64 down-regulated genes for BLCA were identified Supplementary table S16-S18.

### Human tumor samples

Three lung cancer samples were obtained from Shanghai Pulmonary Hospital, and three bladder cancer samples were obtained from Changhai Hospital. This study strictly follows the principles according to the Declaration of Helsinki, with written informed consents obtained from all participants before sample collection according to regular principles. Ethical approvals were obtained from the Institutional Review Board of Changhai Hospital, Second Military University (Approval No. 20191201), and the Medical Ethics Committee of Shanghai Pulmonary Hospital (Approval No. 2024ZD0520206).

### Culture of bladder cancer and lung cancer PTCs

Freshly collected samples were placed in sample preservation medium and transferred to Chengdu OrganoidMed Medical Laboratory on ice to culture PTC models. Briefly, samples were washed with ice-cold phosphate-buffered saline (PBS) supplemented with 10 mM HEPES and 100 U/mL penicillin-streptomycin for 5 times. Necrotic areas and adipose tissue were removed as much as possible. The tissues were minced into small pieces and digested in 5 mL of 1 mM PBS/EDTA containing 200 U/mL of collagenase I for 1 hour. A 40-µm filter was used to collect the dissociated cells. After 10 minutes of centrifugation at 300×g and 4 °C, the cell pellets were re-suspended in PTC growth medium (DMEM/F12 medium supplemented with 5ng/mL msp, 30ng/ mL HGF, 40ng/ mL EGF, 20ng/ mL FGF-basic, 10μM Y-27632 and 1mM N-acetyl-L-cysteine) and seeded at a concentration of 1×105 cells/cm² into an ultra-low attachment plate. The cells were cultured in an incubator at 37 °C with 5% CO₂. The PTC growth medium was refreshed every 2-3 days.

### Drug perturbated PTCs sample preparation and RNA sequencing

PTCs with more than 40 µm in diameter were harvested using 40-µm filter, washed with PBS, and re-suspended with PTC growth medium. Then, 500 μL of the PTC growth medium containing about 400 PTCs was seeded into an ultra-low attachment 24-well plate. Next, 100 μL PTC growth medium containing different compounds was added to the plate, and an equal volume of DMSO as controls. A total of 40 drugs including clinically used chemotherapeutic and targeted drugs were tested, and the information and final concentrations of the compounds were presented in Supplementary table S5. After incubation for 48 hours, PTCs samples were collected and added into Trizol reagent (Invitrogen, CA, USA), and then handed over to OE Biotech Co., Ltd. (Shanghai, China) for the subsequent mRNA library construction and sequencing. Total RNA was extracted using the TransZol Up Plus RNA Kit (TransGen Biotech, China) and the libraries were constructed using VAHTS Universal V10 RNA-seq Library Prep Kit (Premixed Version). RNA integrity was assessed using the Agilent 2100 Bioanalyzer (Agilent Technologies, Santa Clara, CA, USA) and the libraries were sequenced on an Illumina Novaseq 6000 platform and 150 bp paired-end reads were generated. Raw reads of fastq format were firstly processed using fastp and the low quality reads were removed to obtain the clean reads. The clean reads were mapped to the reference genome using HISAT2. FPKM of each gene was calculated and the read counts of each gene were obtained by HTSeq-count.

### UniCure model architecture

UniCure is a foundation model-based framework designed to predict drug-induced transcriptomic perturbations across heterogeneous human cellular systems, from standard cell lines to clinically relevant primary patient samples. Let 𝑋 ∈ ℝ^𝑛^ denote the 𝑛-dimensional gene expression vector representing the transcriptomic state of a cell. The perturbation is defined as a combination 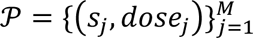 where 𝑠_𝑗_ represents the SMILES representation of the 𝑗-th drug molecule, and *dose*_𝑗_ is the corresponding concentration. The dataset can then be defined as 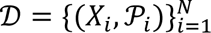, where 𝑋_𝑖_ represents the initial transcriptomic state of the 𝑖 −th cell and 𝒫_𝑖_ denotes the corresponding perturbation applied. The objective of UniCure is to learn a predictive function 𝑓, parameterized by a model, that maps the initial cellular state and perturbation to the post-perturbation transcriptomic state. Formally,

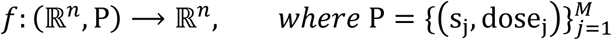

such that

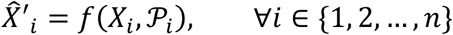

where 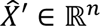 estimates the post-perturbation expression profile 𝑋′. The function 𝑓 is implemented via a foundation model-based architecture comprising four core modules: (1) foundation models with LoRA, (2) FlexPert module, (3) MMD-based alignment, and (4) staged training, as detailed below.

### Drug and Cell Representations with Foundation Models

UniCure derives drug and cellular state embeddings through pre-trained foundation models, formalized as follows:

For a drug molecule represented by its SMILES string 𝑠, UniCure computes its embedding through the pre-trained Uni-mol model ℳ_𝑢𝑛𝑖−𝑚𝑜𝑙_, a 135-layer transformer with 50M parameters. Let Ε_𝑢𝑛𝑖−𝑚𝑜𝑙_ ∈ ℝ^512^, denote the molecular embedding encoding molecule features, obtained by:

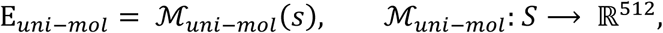

where 𝑆 is the SMILES space. To incorporate dosage effects, the final drug embedding Ε_𝑑𝑟𝑢𝑑𝑑_ is modulated by the logarithmic dose 𝑑𝑜𝑠𝑒:

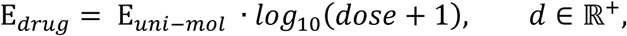

This ensures dose-proportional transcriptional impact while preserving structural semantics. Uni-mol is pre-trained on 2.09 × 10^8^ molecular conformations, extending conventional 1D/2D molecular representation learning (MRL) by explicit 3D geometry modeling.

A cellular state X ∈ ℝ^𝑛^, (gene expression vector for 𝑛 genes) is encoded into a 1280-dimensional embedding Ε_𝑐𝑒𝑙𝑙_∈ ℝ^1280^, via the Universal Cell Embedding (UCE) model ^ℳ^_𝑢𝑐𝑒_:

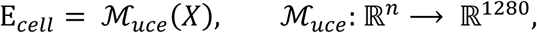

where ℳ_𝑢𝑐𝑒_ processes X by first assigning each gene 𝑘 a sampling probability 𝑝_𝑘_ proportional to its expression level 𝑥_𝑘_:

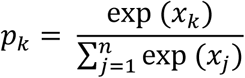

Genes are stochastically selected based on 𝑝_𝑘_, encoded into protein-aware tokens via ESM2 ℳ_𝐸𝑆𝑀2_, and ordered by their chromosomal coordinates. The sorted sequence is then passed through UCE’s 33-layer transformer and decoder layer. The final embedding Ε_𝑐𝑒𝑙𝑙_ is derived by L2-normalizing the CLS token’s output.

### Low-Rank Adaptation (LoRA)

To adapt the UCE model to diverse cellular contexts, spanning cell lines to patient-derived samples, while preserving pre-trained biological knowledge, we implement Low-Rank Adaptation (LoRA). LoRA addresses catastrophic forgetting and computational inefficiency in fine-tuning large models by decomposing weight updates into low-rank matrices^18^. This is particularly important in biological foundation models, where the inherent complexity and stochasticity of biological data make full-parameter fine-tuning more prone to catastrophic forgetting, even when applied to pretrained models. For a pretrained weight matrix W ∈ ℝ^𝑑×𝑘^, LoRA approximates parameter updates as:

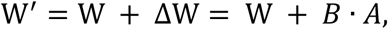

where 𝐴 ∈ ℝ^𝑑×𝑟^ and 𝐵 ∈ ℝ^𝑟×𝑘^ are trainable low-rank matrices (𝑟 = 32 ≪ min(𝑑, 𝑘)), while W remains frozen. During forward propagation, layer outputs are computed as:

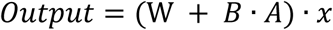

In UniCure, LoRA is applied exclusively to the multi-head attention modules of the last five transformer layers in UCE. This targeted adaptation allows the model to refine context-specific interactions (e.g., tumor microenvironment cues) without overfitting to limited patient data, achieving parameter-efficient customization for individualized predictions.

### FlexPert Module

To flexibly accommodate multi-drug inputs of arbitrary number and order, the FlexPert module was designed to overcome the limitations of conventional models that require fixed-dimensional drug embeddings. Inspired by attention-based architectures in natural language processing^40, 41^, which handle variable-length token sequences, our design treats both drug and cell embeddings as ordered sequences after initial representation learning. While raw gene expression vectors are inherently unordered—since gene positions are interchangeable without altering cellular identity—the embedding process by the UCE induces a semantic structure, enabling meaningful spatial locality. We therefore apply a sliding window mechanism to generate overlapping segments (akin to tokens) from both drug and cell embeddings. These windowed segments are then fed into a cross-attention mechanism, where cell-derived windows serve as queries and drug-derived windows as keys and values. This setup allows the model to simulate perturbation effects by integrating drug information directly into the cellular embedding in a targeted, biologically meaningful manner. Compared to traditional MLP-based encoders or decoders, the attention formulation offers greater expressiveness and aligns more closely with the mechanistic notion of drug-induced perturbation.

Given a cell embedding Ε ∈ ℝ^1280^ and one or more drug embeddings 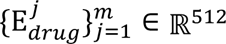, the module operates as follows:

1. Sliding Window Processing: Each embedding is reshaped into a 2D tensor via sliding windows (window size = 32, stride = 16.) For the cell embedding:

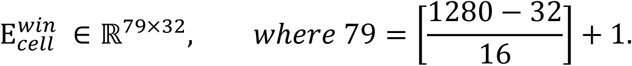 For each drug embedding 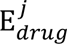:

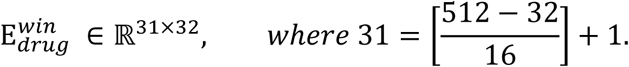 Multi-drug embeddings are concatenated vertically:

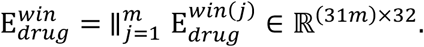
2. Cross-Attention Mapping: The cell and drug window tensors are projected into query, key, and value spaces via single-layer linear networks:

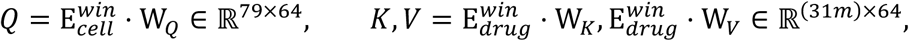

where W_𝑄_, W_𝐾_, W_𝑉_ ∈ ℝ^32×64^ are learnable weights.
3. Attention Computation: The scaled dot-product attention is computed as:

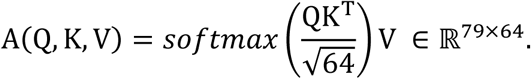 The output is flattened into a 1D vector 𝑧 ∈ ℝ^5056^.
4. FlexPert Decoder: A multi-layer perceptron (MLP) with ReLU activation transforms 𝑧:

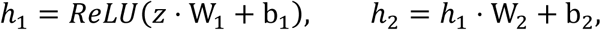

where W_1_, W_2_, W_3_ ∈ ℝ^5056×5056^.
5. UniCure Decoder: The final perturbed transcriptomic state 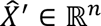 is generated by the UniCure decoder:

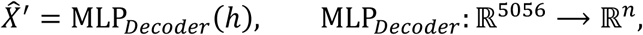

with a hidden layer of 2048 units and ReLU activation.

### Maximum Mean Discrepancy Module

To address the challenge of unpaired pre- and post-perturbation data, UniCure employs a Maximum Mean Discrepancy (MMD) loss with an additional transport cost constraint. This design reflects a fundamental limitation of biological sequencing technologies: the destructive nature of single-cell profiling prevents precise one-to-one pairing of cellular states before and after drug treatment. Prior transcriptome-based models have addressed this issue through suboptimal strategies, including (1) averaging cells under identical conditions, which sacrifices cellular heterogeneity and reduces biological resolution^8, 11^; (2) randomly pairing cells from the same condition, which introduces noise and inconsistency^10, 12^; or (3) applying optimal transport algorithms to align distributions, which cannot generalize to unseen compounds^29^. In contrast, UniCure’s formulation enables distribution-level alignment without relying on cell-level pairing, while maintaining the capacity to generalize across novel drug perturbations:

1. MMD Definition: The MMD distance between the predicted post-perturbation distribution 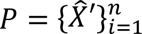 and the observed post-perturbation 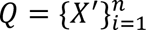, under the same perturbation conditions (cell line, experimental protocol, drug dose, and compound identity), is computed using a RBF kernel 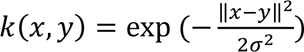, where 𝜎 = 2.0 is the kernel bandwidth. The MMD loss is defined as:

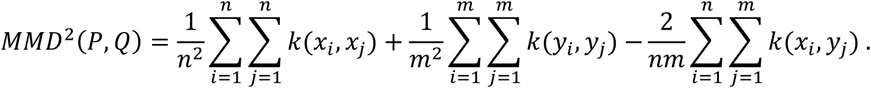
2. Transport Cost Constraint: To regularize the perturbation magnitude, we introduce a Euclidean distance penalty between the original unperturbed state X_𝑖_ and the predicted perturbed state 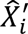. The constraint term is scaled by a coefficient 𝜆 = 0.01:

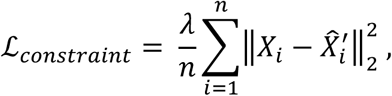 The Euclidean penalty ensures predicted perturbations remain biologically plausible by limiting deviations from the original state. The total loss function is defined as:

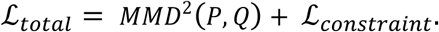

### Staged Training Strategy

UniCure adopts a biologically inspired two-phase training protocol to decouple cellular context learning from perturbation modeling, enhancing computational efficiency and functional modularity.

#### Phase 1: Cellular Context Stabilization

In this phase, the model is trained on unperturbed cell states to establish robust baseline representations. The following components are activated:

- UCE-Encoder with LoRA: Low-rank adaptation layers fine-tune the last five transformer layers of UCE.
- UCE-Decoder: Reconstructs baseline transcriptomic profiles.
- Query Projection (W_𝑄_): Maps sliding-window-processed cell embeddings to query vectors.
- UniCure-Decoder: Predicts gene expressions.

Trainable Parameters:

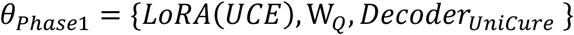

Objective: Minimize the mean squared error (MSE) between predicted and observed unperturbed states:

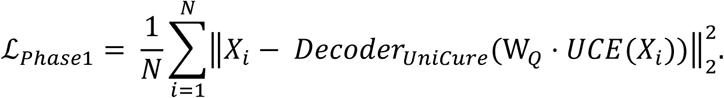

#### Phase 2: Perturbation Modeling

Building on Phase 1, this phase trains the model on drug-perturbed cell states with the following updates:

- Frozen Components: All parameters from Phase 1 remain fixed.
- Activated Modules:

- Key/Value Projections (W_𝐾_, W_𝑉_): Map drug embeddings to key/value vectors.
- FlexPert Decoder: Transforms cross-attention outputs into perturbation-aware embeddings.

Trainable Parameters:

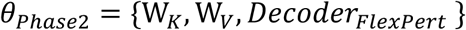

Objective: Minimize the combined MMD and transport constraint loss between predicted and observed perturbed states:

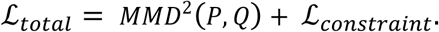

Phase 1 stabilizes cellular context encoding without drug interference. Phase 2 isolates perturbation learning, mimicking biological systems which baseline states precondition drug responses. This staged strategy ensures that UniCure hierarchically learns cellular dynamics, aligning with both biological principles and computational scalability requirements.

### Transfer Learning for Single-Cell and PTC Datasets

To enable UniCure’s generalization beyond bulk training data, we implemented tailored fine-tuning strategies for single-cell datasets, based on the model pretrained on LINCS2020.

- Single-cell adaptation (SciPlex3 and SciPlex4) For adaptation to single-cell data, we followed a transfer learning strategy inspired by chemCPA^29^, applying two key modifications:

- Encoder Replacement: The LoRA-adapted UCE encoder was replaced with the native UCE encoder, as the former was optimized for bulk transcriptomes. The native UCE encoder is better suited to retain the cell-to-cell variability and structure intrinsic to single-cell expression data.
- Output Layer Extension: To support the broader gene set in single-cell datasets (𝐺_𝑠𝑐_ = 1923), we augmented the UniCure decoder 𝑔_𝜃_ with a task-specific linear layer W ∈ ℝ^𝐺𝑠𝑐 ×𝑑^, yielding final prediction:

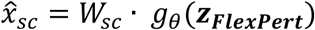

Where 𝑧_𝑓𝑢𝑠𝑖𝑜𝑛_ is the cross-modality embedding output from the FlexPert module. The FlexPert module and W_𝑠𝑐_ were fine-tuned end-to-end using single-cell perturbation data. The SciPlex4 model was initialized from the SciPlex3-trained checkpoint and updated using the same transfer strategy.
- PTC fine-tuning strategy To align the decoder’s output to the 978-dimensional landmark gene space 𝐺_𝑝𝑡𝑐_ = 978, a linear layer W_𝑝𝑡𝑐_ ∈ ℝ^978×𝑑^ was appended:

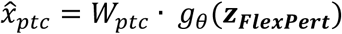 During PTC fine-tuning, only W_𝑝𝑡𝑐_ was updated:

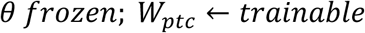 This lightweight adaptation enabled effective tuning on limited patient data while preserving core biological priors encoded by the pretrained UniCure model.

### Personalized Drug Prioritization Strategy

To prioritize therapeutic candidates for individual patients, we designed a UniCure-based screening strategy that leverages paired tumor–normal transcriptomes. Let 𝑥^𝑡𝑢𝑚𝑜𝑟^ ∈ ℝ^𝐺^ and 𝑥^𝑛𝑜𝑟𝑚𝑎𝑙^ ∈ ℝ^𝐺^ denote the gene expression profiles of tumor and adjacent normal tissues for a given patient, where 𝐺 is the number of genes (e.g., 𝐺=978 landmark genes). For each drug 𝑑 ∈ 𝓓 in the compound library 𝓓, UniCure predicts the perturbed tumor transcriptome 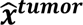.

We compute the drug-induced differential expression vector as:

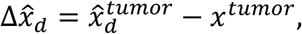

and the disease-associated differential expression as

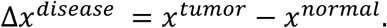

To quantify the potential of each drug 𝑑 to reverse tumor-specific transcriptional changes, we calculate the Pearson correlation coefficient:

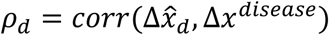

A lower value of 𝜌_𝑑_ suggests stronger antagonism between the predicted drug response and the disease signature, indicating a higher potential to therapeutically revert the tumor phenotype. All drugs are thus ranked in ascending order of 𝜌_𝑑_ to generate a patient-specific prioritization list. This ranked vector can also serve as a new patient-level feature representation for downstream analysis such as clustering or treatment outcome prediction.

### Screening candidates based on gene signatures

Building upon the computational framework of the CMap connectivity score^1, 2^, UniCure adopts a similarity approach for therapeutic candidate screening.

Step 1: Given the SMILES of the screening compounds library and unperturbed transcriptional profiles of specific cell lines, UniCure predicts the perturbed transcriptional profiles of all compounds and generate compounds signature are by differential expression analysis.

Step 2: Disease signatures representing the disease states are calculated and genes are ranked based on their fold-change values, which include an up-regulated gene set and a down-regulated gene set.

Step 3: The CMap connectivity score method is employed to calculate enrichment scores, which measure the connectivity of screening compound profiles with the disease signatures.

3.1. Given a ranked list of drug genes signature L with n genes and a disease signature S with m genes.

3.2. Given a vector V indicating the position (1, 2, …, n) of each gene in L and sort the genes in S in ascending order such that V(i) is the position of gene i, where i =1, 2, …, t. Compute the following two values:

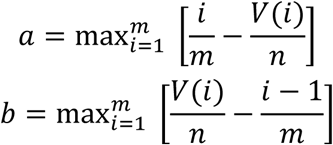

And hence the KS score is defined as:

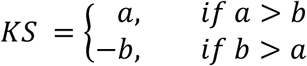

L refers to the gene list ranked by gene expression of compound, S refers to the up-regulated or down-regulated genes in disease signature.

If sign(KS_up_) = sign(KS_down_), CMAP score = 0, otherwise, CMAP score = KS_up_ - KS_down_. Finally, connectivity scores were normalized to range from −1 to 1, and compounds were subsequently ranked according to these scores. The top-ranked compounds were then recommended as potential therapeutic candidates for relevant diseases.

### Cell culture

MDA-MB-231 cells and A549 cells were obtained from Shanghai Honsun Biological Technology Co., Ltd, 5637 cells were obtained from Shanghai BinSui Biological Technology Co., Ltd. All the natural products in the experiment were purchased from TargetMol Chemicals Inc (Shanghai, China). MDA-MB-231 and A549 cells were cultured in DMEM supplemented with 10% fetal bovine serum (FBS) and 1% streptomycin and penicillin (P/S), while 5637 cells were cultured in RPMI 1640 supplemented with 10% FBS and 1% P/S. The above-mentioned cell lines were all cultured in an incubator at 37 ° C and 5% CO2.

### Cell Viability Assay

Cells were counted and seeded in 96-well plates at a density of 5000 cells per well. The following day, cells were treated with compounds at predefined concentrations. After incubation for 48 hours, cell viability was assessed using an CCK-8 kit, and dose-response curves were generated using GraphPad Prism 9.5.

### Colony Formation Assay

Cells were counted and seeded in 12-well plates (2000 MDA-MB-231 cells, 1000 A549 cells or 2000 5637 cells per well). The following day, cells were treated with compounds at predefined concentrations. After 2 weeks, cells were fixed with 4% paraformaldehyde and then stained with Crystal Violet Staining Solution (C0121, Beyotime). Colonies containing ≥50 cells were counted manually under an inverted microscope.

### Bioinformatics Analyses and Visualization

Pathway enrichment analyses were conducted using the R package clusterProfiler^42^ (v4.6.2) to identify biological pathways associated with differentially expressed genes. Survival analyses, including Kaplan–Meier estimation and Cox proportional hazards modeling, were performed using survival (v3.8-3) and survminer (v0.5.0) packages in R.

Dimensionality reduction (t-SNE), K-means clustering, and correlation density visualizations were implemented in Python (v3.10.13) using standard scientific computing libraries. All other visualizations, including heatmaps, bar plots, and bubble charts, were generated in R (v4.2.1).

## Supporting information

Extend Data Figures 1-9

## Data availability

Unpublished TNBC PTC data were shared by a collaborating clinical team under agreement. These data are not publicly available due to ongoing analysis and restrictions related to patient privacy, but may be available upon reasonable request with the consent of the original investigators. Publicly available clinical transcriptomic data were obtained from the UCSC Xena Browser (https://xenabrowser.net). The raw expanded CMap LINCS Resource 2020 is available at https://clue.io/data/CMap2020#LINCS2020. The raw sci-Plex3 and sci-Plex4 dataset is available at http://www.perturbase.cn/, and the raw LUAD and BLCA cohort of the TCGA is available at https://xenabrowser.net/. The analysis and results data generated in this study are provided in the Source Data file. All data are available from the corresponding author upon request. Source data are provided with this paper. High-throughput data of Drug perturbated PTCs sample can be accessed at ZexiChen502/UniCure.

## Code availability

The code of UniCure is freely available at ZexiChen502/UniCure.

## Acknowledgements

This work has been supported by National Key R&D Program of China (2022YFA1004800, 2022YFC3502000, 2025YFF1207900), Natural Science Foundation of China (82430119, T2341007, T2350003, 12131020, 42450084, 42450135, 12326614, and 12426310), Science and Technology Commission of Shanghai Municipality (23JS1401300), the ability establishment of sustainable use for valuable Chinese medicine resources (2060302), Chinese Academy of Medical Sciences (CAMS) Innovation Fund for Medical Sciences (2023-I2M-3-009), Zhejiang Province Vanguard Goose-Leading Initiative (2025C01114), Hangzhou Institute for advanced study of UCAS (2024HIAS-P004), the Chenguang Program of Shanghai Education Development Foundation and Shanghai Municipal Education Commission (23CGA45, Saisai Tian), and JST Moonshot R&D (JPMJMS2021).

