## Supplementary material for "UniCure: A Foundation Model for Predicting Personalized Cancer Therapy Response": Extend Data Figures 1-9

Extended Data Fig. 1 Training and Validation Loss and R² Metrics for UCE LoRA Pre-Training and SciPlex Datasets

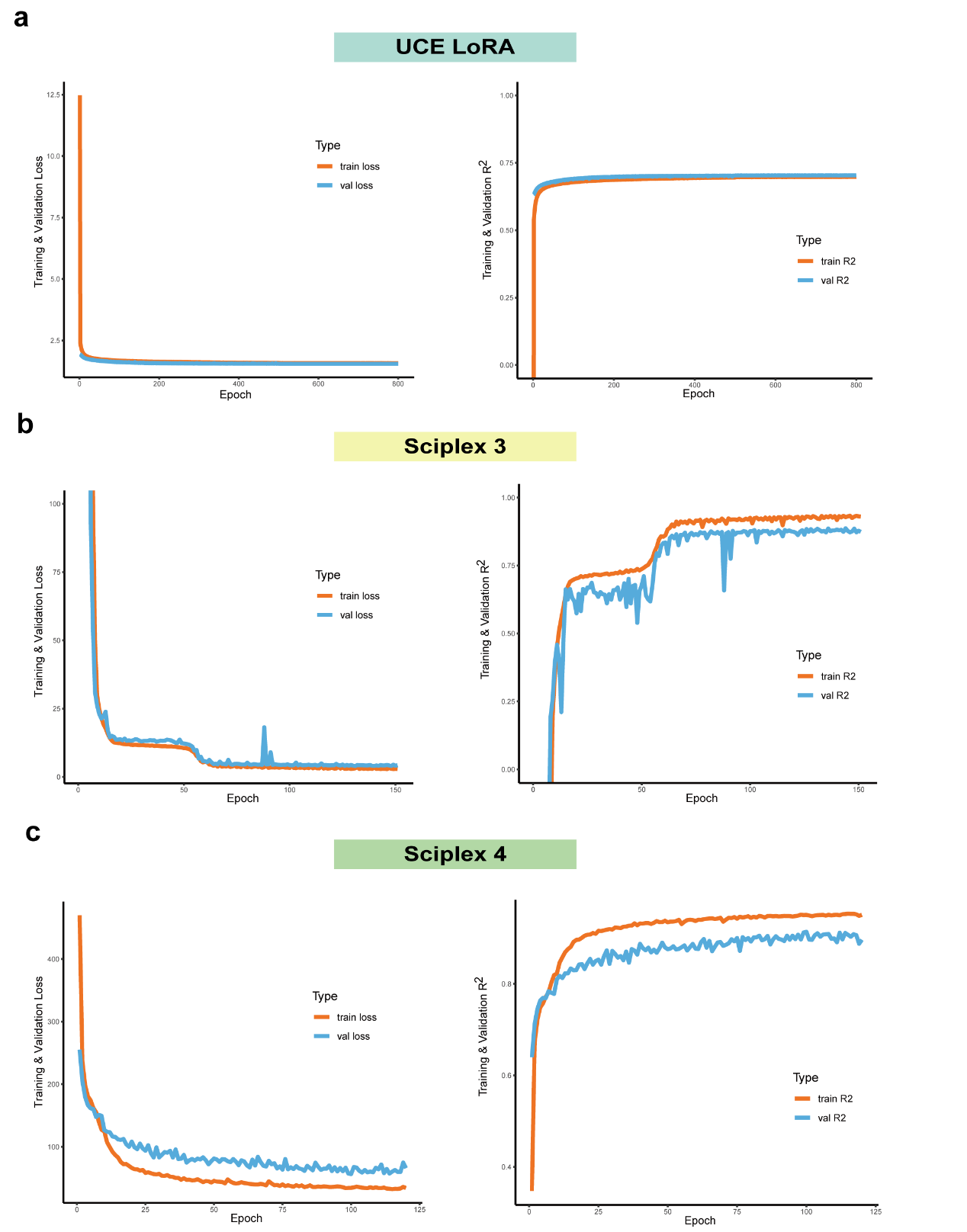

**a**, UCE LoRA pre-training shows a stable decline in both training and validation loss across epochs, coupled with consistent improvements in training and validation R² metrics, indicating effective learning and generalization. **b-c**, Training on the SciPlex 3 (**b**) and SciPlex 4 (**c**) dataset reveals consistent convergence patterns, with validation R² steadily improving over epochs, confirming the robustness and adaptability of UniCure across different datasets and perturbation contexts.

Extended Data Fig. 2 t-SNE Visualization of UniCure Predictions and Real Perturbation Profiles in SciPlex3 Dataset

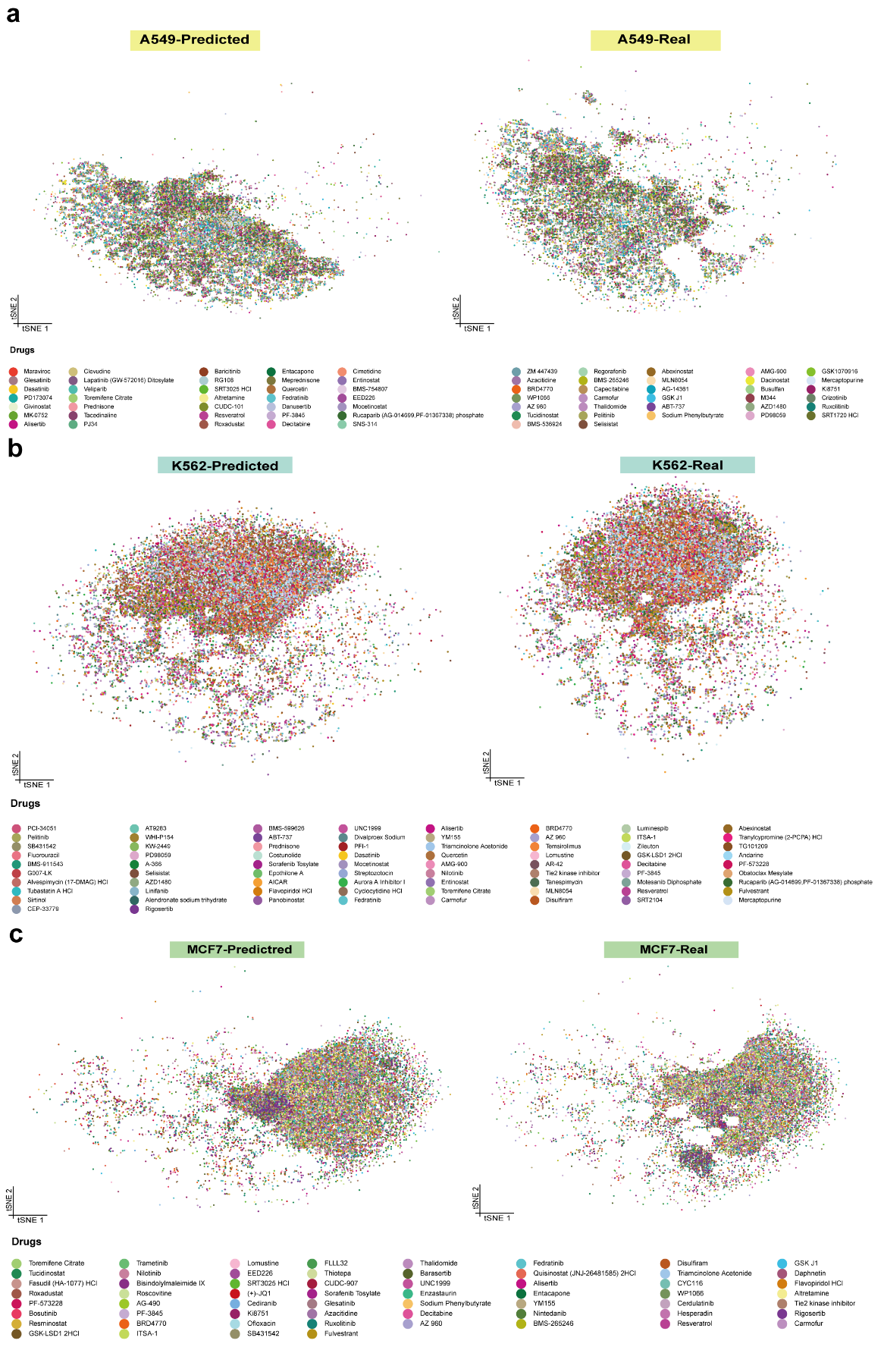

**a-c**, t-SNE projections comparing UniCure’s predicted transcriptomic profiles with real perturbation profiles from the SciPlex3 single-cell dataset across three cell lines: (**a**) A549, (**b**) K562, and (**c**) MCF7. Predicted perturbation profiles (left) closely align with real data distributions (right), preserving the global cell-type structure and recapitulating the overall latent space geometry.

Extended Data Fig. 3: UniCure-Predicted and Experimentally Observed Gene Expression Profiles in MCF7 and PC3 Cells

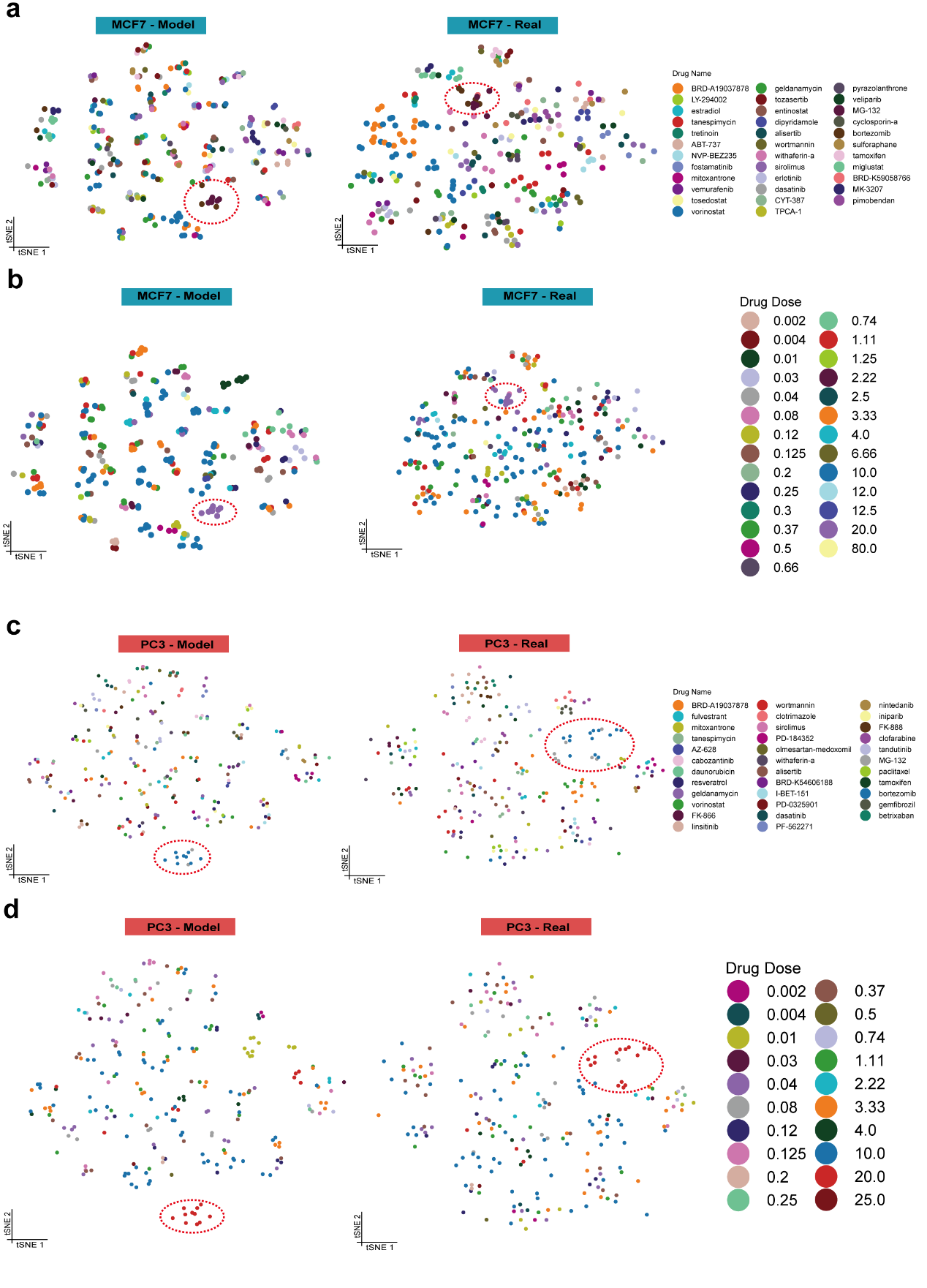

**a-b**, UniCure-predicted (left) and experimentally observed (right) gene expression profiles in MCF7 cells, colored by compound identity (**a**) and drug dose (**b**). **c-d**, UniCure-predicted (left) and experimentally observed (right) gene expression profiles in PC3 cells, colored by compound identity (**c**) and drug dose (**d**).

Extended Data Fig. 4: Comparison of Predicted and Observed Target Gene Expression Across Colon, Lung, and Prostate Adenocarcinoma Cell Lines

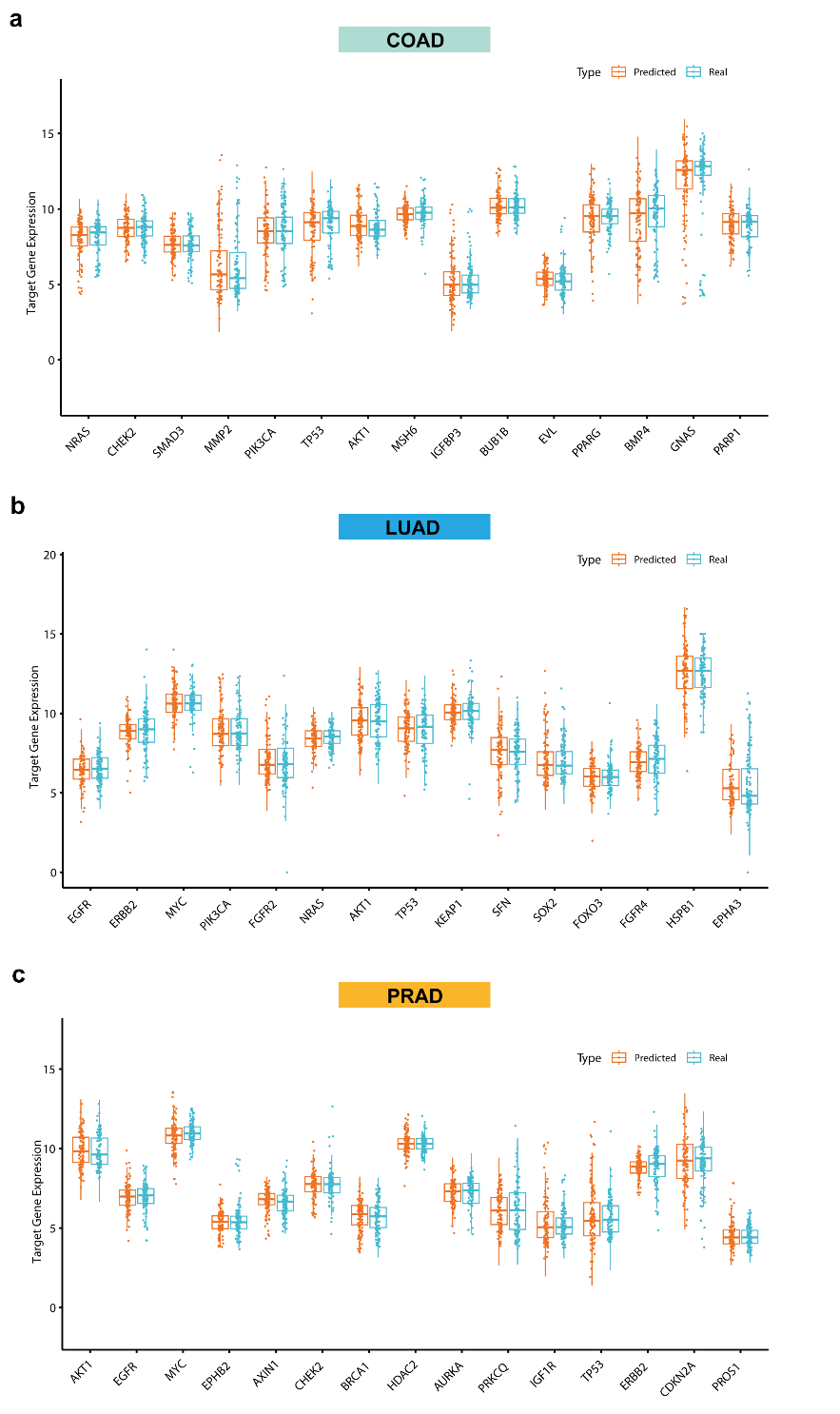

**a-c**, Comparison between UniCure-predicted and experimentally observed expression of target genes across all colon adenocarcinoma (**a**), lung adenocarcinoma (**b**) and prostate adenocarcinoma (**c**) cell lines in the LINCS2020 dataset.

Extended Data Fig. 5: Volcano Plots of DEGs Predicted by UniCure in MCF7 Cells Treated with PI3K Inhibitors

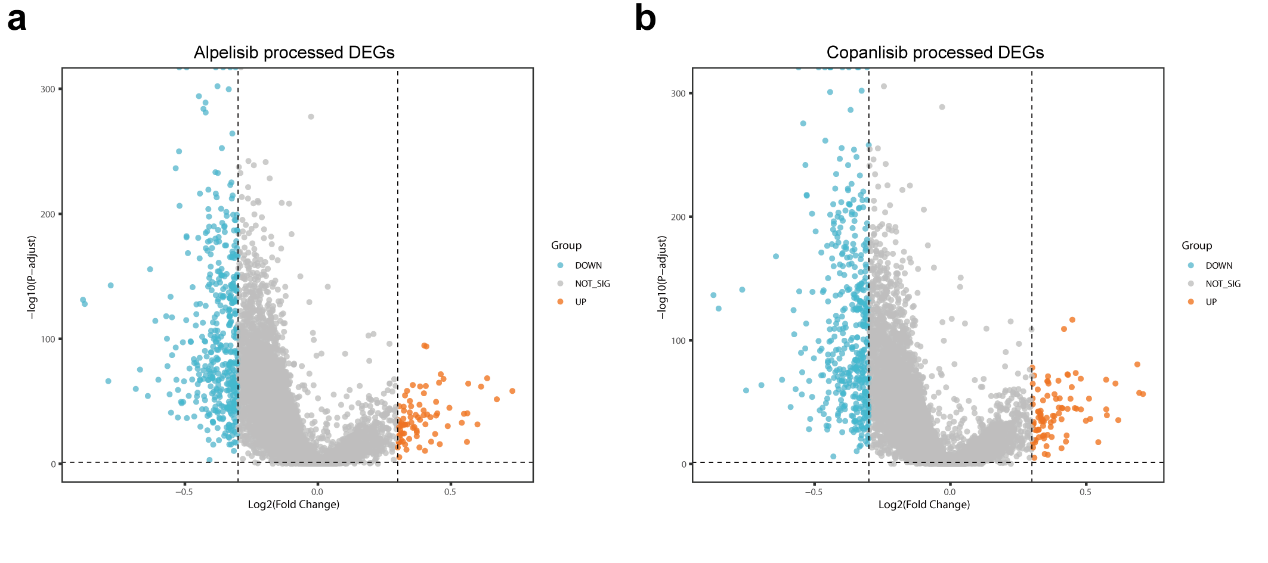

**a-b**, Volcano plots illustrating UniCure's predictions of differentially expressed genes (DEGs) in response to PI3K inhibitors including Alpelisib (**a**) and Copanlisib (**b**) in the MCF7 cell line, which harbors a PIK3CA mutation.

Extended Data Fig. 6: Microscopy, Histology, and Immunofluorescence Characterization of Bladder and Lung Cancer PTC

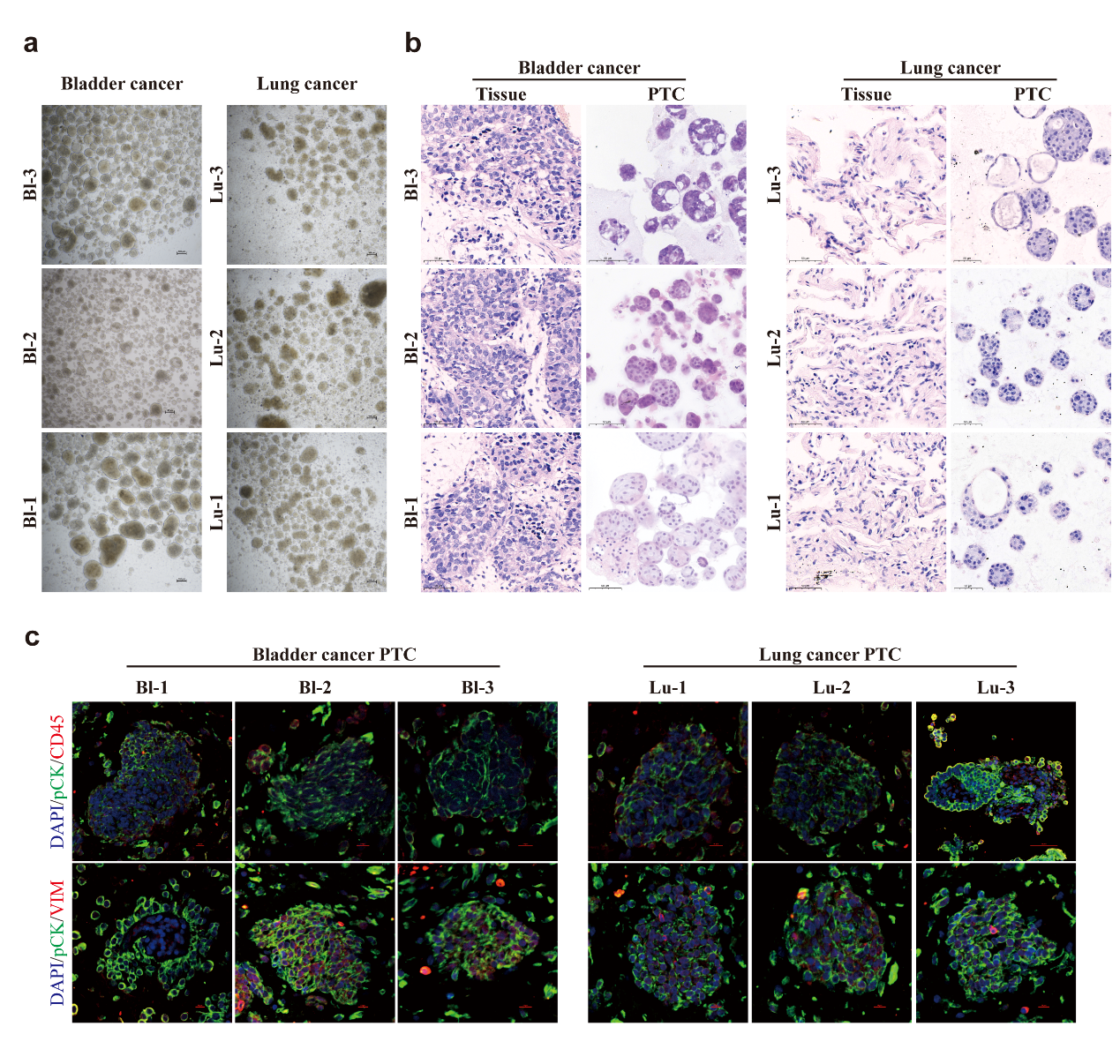

**a**, Microscopic images showing three-dimensional self-assembly of bladder (Bl1-3) and lung (Lu1-3) cancer PTCs cultured in Matrigel-free conditions. **b**, Hematoxylin and eosin (H&E) staining of surgically resected parental tumor tissues and corresponding PTCs demonstrate morphological fidelity. **c**, Immunofluorescence staining of patient-derived tumor-like cell clusters (PTCs) from bladder (Bl1-3) and lung (Lu1-3) cancer samples.

Extended Data Fig. 7: Performance Evaluation of UniCure on PTC Test Data and Real-World Patient Samples Across Cancer Types

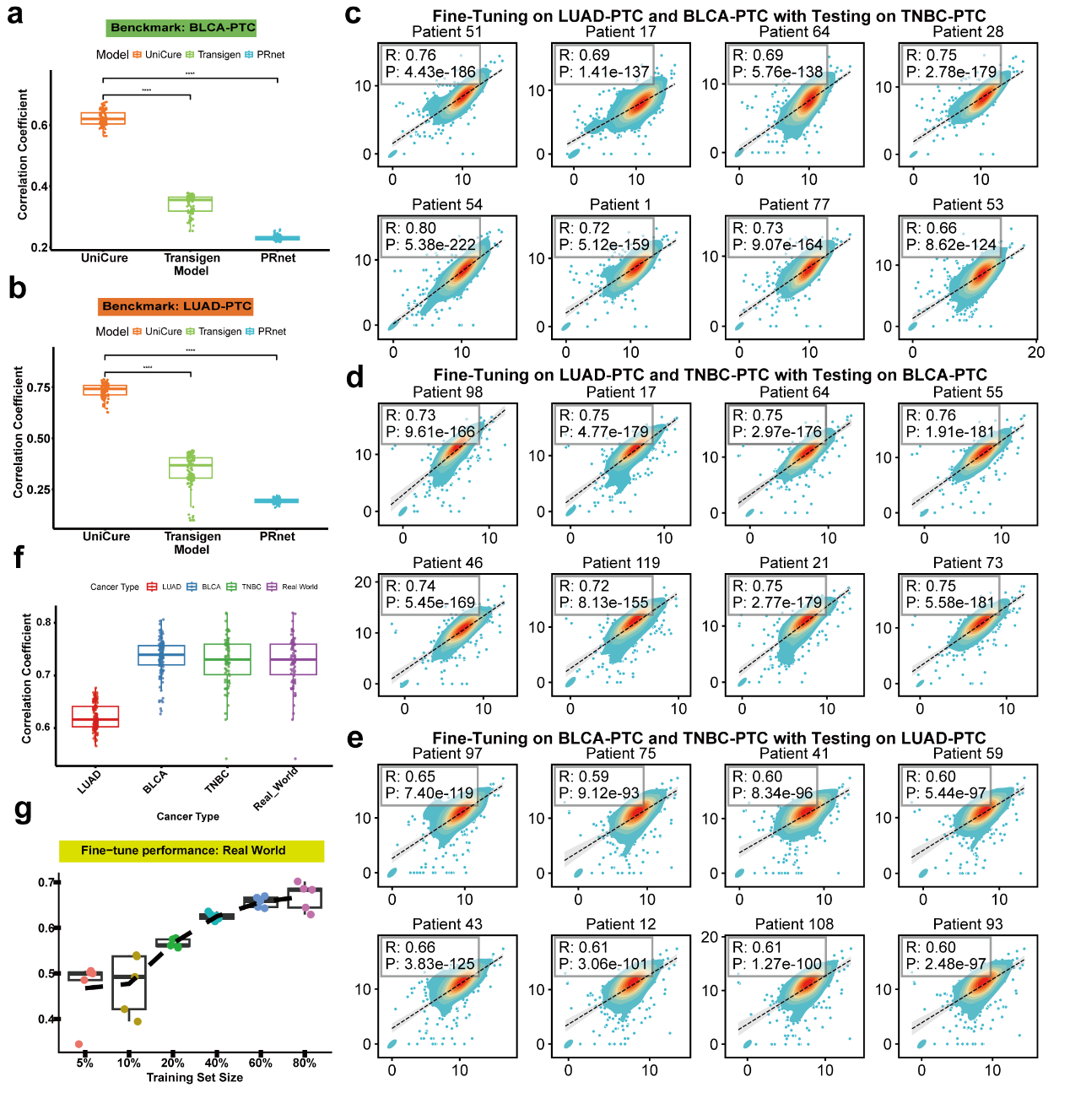

**a-b**, Benchmark comparison of UniCure, TransiGen, and PRnet on BLCA-PTC (**a**) and LUAD-PTC (**b**) test data. Wilcoxon rank-sum test, **** p < 0.0001. **c-e**, Scatter and density plots comparing predicted versus true gene expression profiles for 8 randomly selected real-world patient samples. **c**, Predictions tested on TNBC-PTC after fine-tuning on LUAD-PTC and BLCA-PTC. **d**, Predictions tested on BLCA-PTC after fine-tuning on LUAD-PTC and TNBC-PTC. **e**, Predictions tested on LUAD-PTC after fine-tuning on BLCA-PTC and TNBC-PTC. **f**, Predictive performance after fine-tuning, measured by Pearson correlation coefficients: Predictions tested on TNBC-PTC after fine-tuning on LUAD-PTC and BLCA-PTC; Predictions tested on BLCA-PTC after fine-tuning on LUAD-PTC and TNBC-PTC; Predictions tested on LUAD-PTC after fine-tuning on BLCA-PTC and TNBC-PTC; Predictions tested on real-world patient data after fine-tuning on LUAD-PTC, BLCA-PTC, and TNBC-PTC. **g**, Fine-tuning performance of UniCure exclusively on real-world patient data, illustrating the relationship between training set size and predictive accuracy.

Extended Data Fig. 8: Extended data from UniCure validation in a public clinical cohort

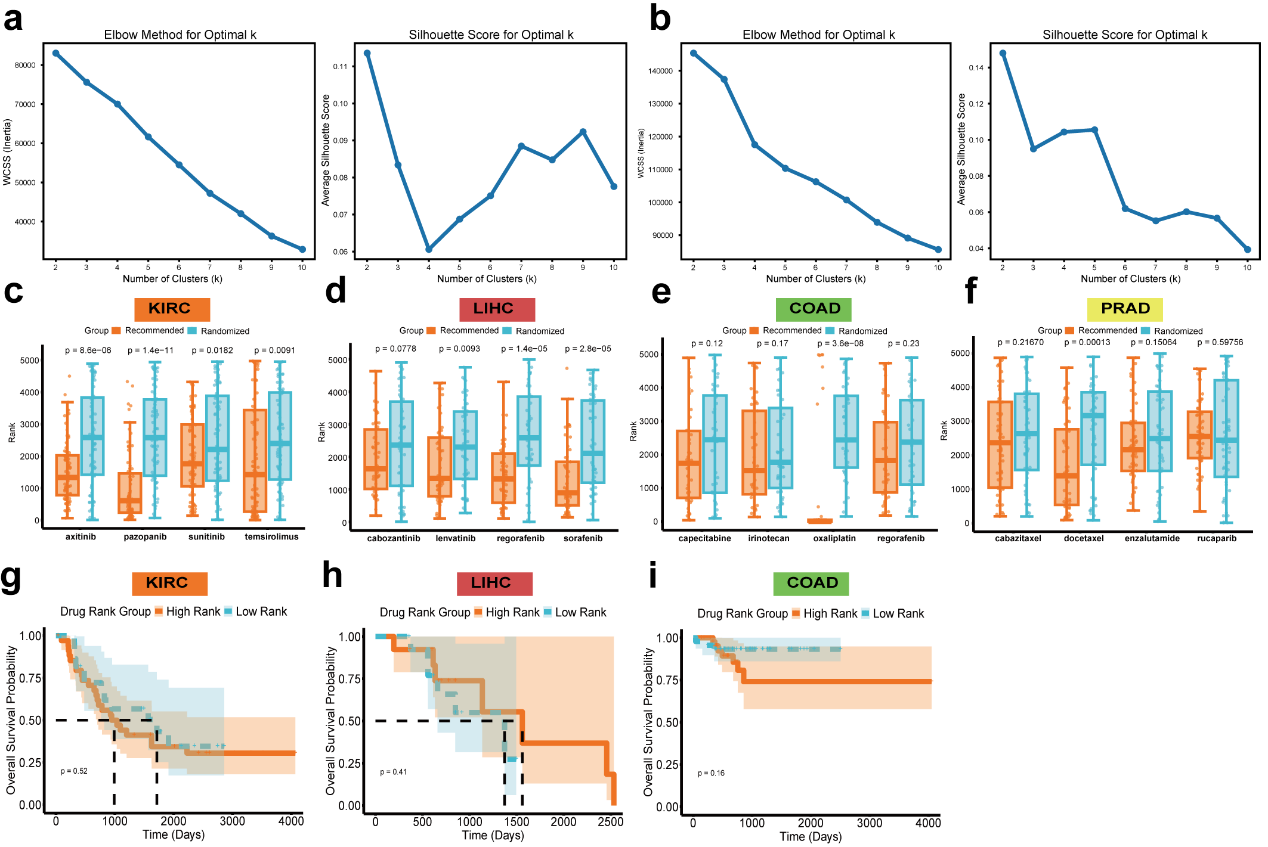

**b**, Determination of optimal cluster number (k) for K-means clustering applied to personalized drug ranking vectors by within-cluster sum of squares (WCSS, left) and the silhouette score analysis (right) metrics in late-stage breast cancer (BRCA, **a**) and clear cell renal cell carcinoma (KIRC, **b**) cohorts. **c-f**, Predicted rank distributions of clinically relevant therapies across public cohorts of KIRC (**c**), LIHC (**d**), COAD (**e**), and PRAD (**f**) patients, compared to randomized controls; significance assessed by Wilcoxon rank-sum test. **g-i**, Kaplan–Meier survival analysis for COAD (**g**), KIRC (**h**), and LIHC (**i**) patients, stratified by whether the administered drug ranked in the top 30% (High Rank) or bottom 70% (Low Rank) of UniCure’s predicted rankings.

Extended Data Fig. 9: ROC Curves Evaluating Classification Performance of Gene Expression Signatures in TNBC, LUAD, and BLCA

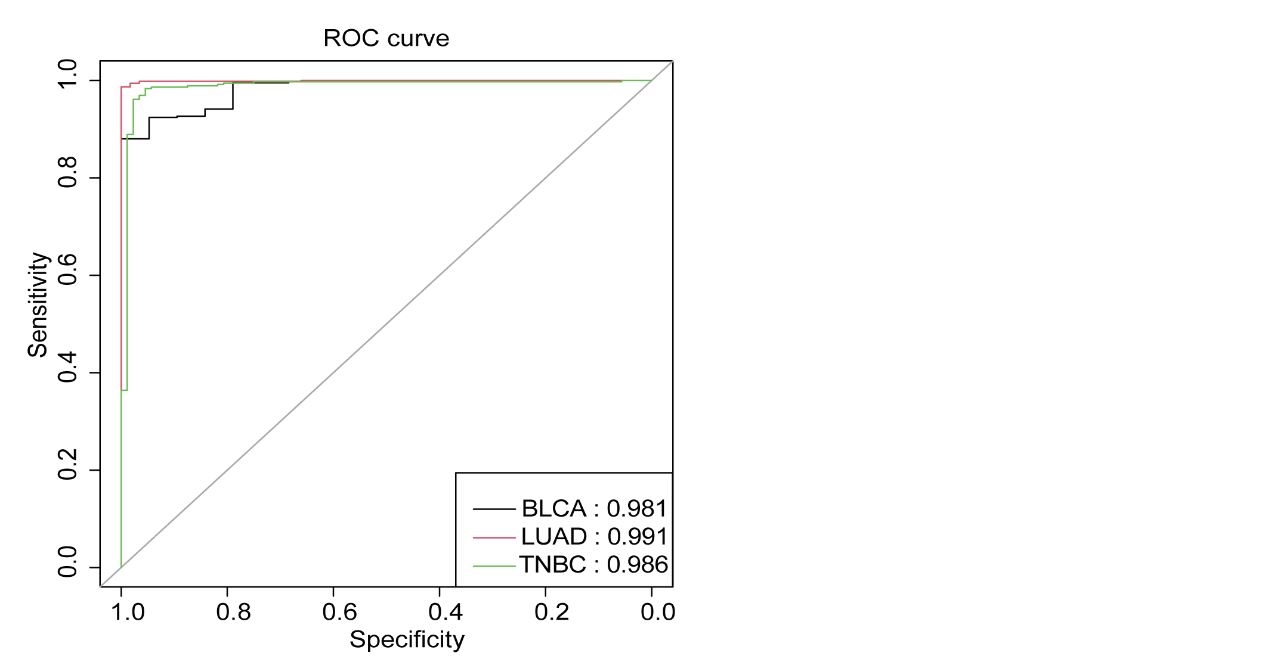

Receiver operating characteristic (ROC) curves evaluating the classification performance of disease gene expression signatures for TNBC, LUAD, and BLCA in distinguishing tumor samples from adjacent normal tissue samples.
