## Supplemental Method, Supplemental Figure 1 for "UniCure: A Foundation Model for Predicting Personalized Cancer Therapy Response"

Supplementary Methods

**Evaluation Metrics**

Model performance is quantified on the held-out test set using three metrics:

1. R-squared (R^2^): Measures the proportion of variance in observed transcriptomic changes explained by predictions:

$R^{2}=1-\frac{\sum_{i=1}^{N} \left( y_{i}-\hat{y}_{i} \right)^{2}}{\sum_{i=1}^{N} \left( y_{i}-\bar{y}_{i} \right)^{2}}$ ,

where $y_{i}$ is the observed expression, $\hat{y}_{i}$ the predicted value, and $\bar{y}_{i}$ the observed mean.

2. Pearson Correlation Coefficient: Evaluates linear correlation between predicted and observed expression values:

$r= \frac{\sum_{i=1}^{N} (y_{i}-\bar{y})(\hat{y}_{i}-\bar{\hat{y}})}{\sqrt{\sum_{i=1}^{N} {(y_{i}-\bar{y})}^{2}} \sqrt{\sum_{i=1}^{N} {(\hat{y}_{i}-\bar{\hat{y}})}^{2}}}$ .

3. Spearman’s Rank Correlation: Assesses monotonic relationships by ranking predictions and observations:

$\rho=1-\frac{6\sum_{i=1}^{N} d_{i}^{2}}{N(N^{2}-1)}$ ,

where $d_{i}$ is the rank difference for the $i$-th sample.

These metrics collectively capture both linear and non-linear associations, ensuring robust assessment of transcriptomic perturbation accuracy.

**Computational Details**

Due to constraints in computational resources, comprehensive hyperparameter optimization was not conducted for the current model. The training protocol is implemented with the following configurations :

Phase 1 (Cellular Context Stabilization):

- Epochs: 800, with an initial learning rate of $1\times{10}^{-4}.$
- Learning Rate Schedule: A multi-step scheduler reduces the learning rate by a factor of 10 at epoch 400.
- Batch Size: Fixed at 64, with samples randomly shuffled across all unperturbed cells.

Phase 2 (Perturbation Modeling):

- Early Stopping: Training terminates if the validation loss does not improve for 20 consecutive epochs.
- Learning Rate: Initialized at $1\times{10}^{-5}.$
- Dynamic Batch Size: Samples are grouped by perturbation conditions (cell line, experimental protocol, drug dose, and compound identity), resulting in variable batch sizes to ensure intra-batch condition consistency.
- Data Split: To prevent data leakage and ensure a rigorous evaluation, samples were first grouped based on shared perturbation conditions. These groups were then randomly split into training, validation, and test sets (approximately 8:1:1), guaranteeing that samples from the same experimental condition were confined to a single dataset (training, validation, or test).

Hardware and Framework:

- The model is trained on 8 NVIDIA A100 GPUs using the Accelerate library with DeepSpeed optimization for distributed parallel computing.
- Phase 1 converges within 3 days, while Phase 2 requires 7 days due to increased complexity from perturbation-aware training and dynamic batching.

Model training and evaluation were conducted using PyTorch (v2.5.1) with CUDA version 12.2 for GPU acceleration.

Supplementary Figures

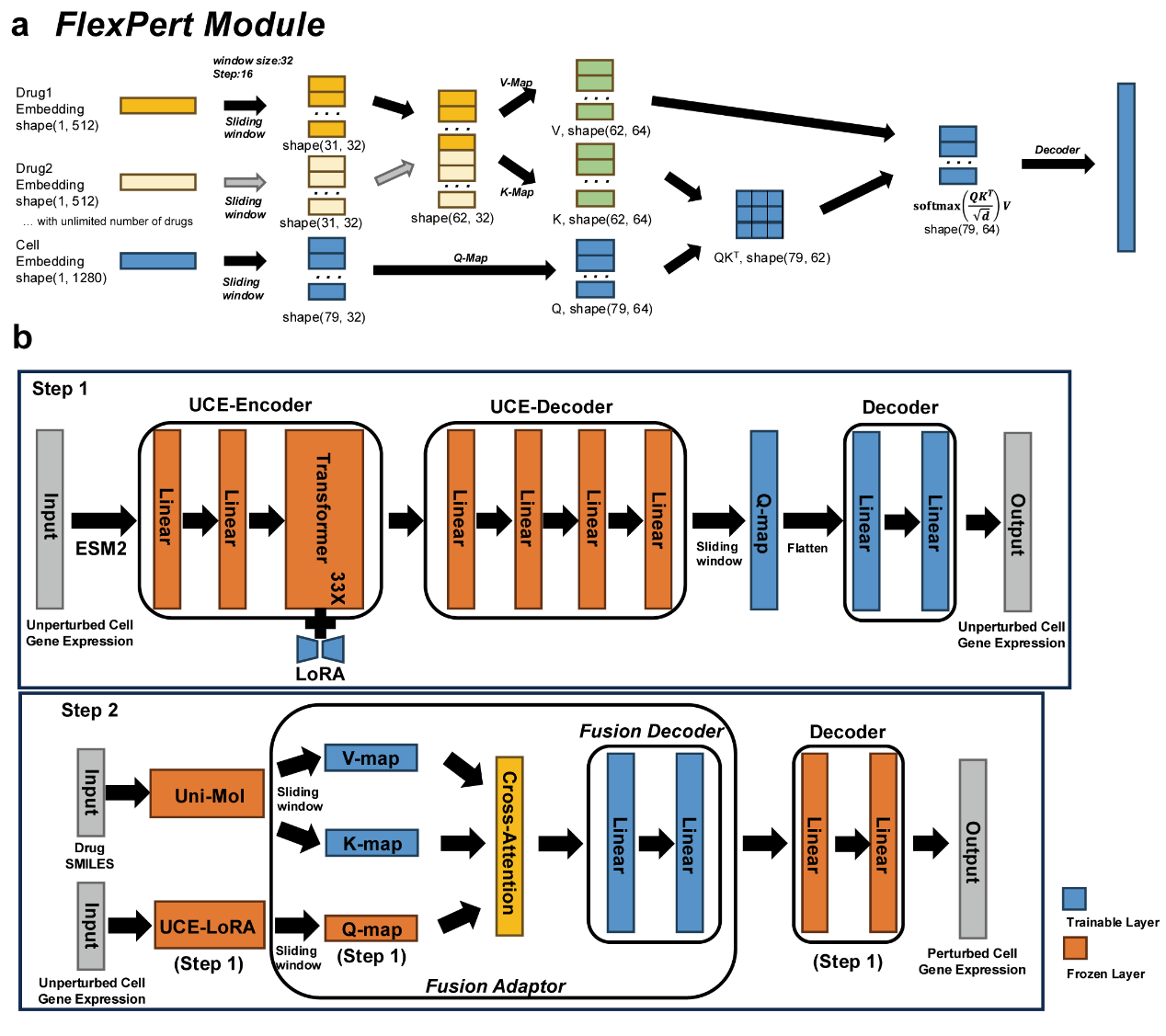

**Supplementary Figure 1 FlexPert module details and model architecture a**, FlexPert module integrates drug and cell embeddings using sliding window-enhanced cross-attention mechanisms. **b**, Overview of the UniCure architecture and staged training strategy.
